# Multiscale Modelling of Multiple Sclerosis Initiation and Progression

**DOI:** 10.64898/2026.01.04.697229

**Authors:** Thomas Hillen, Adrianne L. Jenner

## Abstract

Multiple Sclerosis (MS) is an autoimmune disease that affects the central nervous system. It can lead to inflammation, neurodegeneration, and physical or cognitive disability. Currently, no cure for MS exists, but medications are available to slow its progression. To date, mathematical modelling of MS has focussed on a few aspects of the disease, but an overall modelling framework is missing. In this paper, we propose a multiscale paradigm for the mathematical modelling of MS. Based on the underlying biological scale, we propose six consecutive modelling levels and develop the first three model levels in this work using systems of ordinary differential equations. We test if these models can describe known effects related to MS disease risk, with particular focus on estrogen, vitamin D, Epstein-Barr virus (EBV) and HLA-DR mutations. We first show that periodic disease outbreaks are possible in this framework through interactions by antigen-presenting cells, regulatory cells and memory B cells. We show that the presence of Epstein-Barr virus infections can initiate the disease, low and high levels of estrogen and vitamin D deficiency can alleviate it, mutations in the HLA-DR gene can promote MS. We can identify a stable steady state of our model as the widely discussed MS prodrome, and we find that memory B-cells play a dominant role in the disease progression. We hope that this framework may serve as a reference for the development and comparative evaluation of future mathematical and computational models of MS.

**Graphical Abstract:** 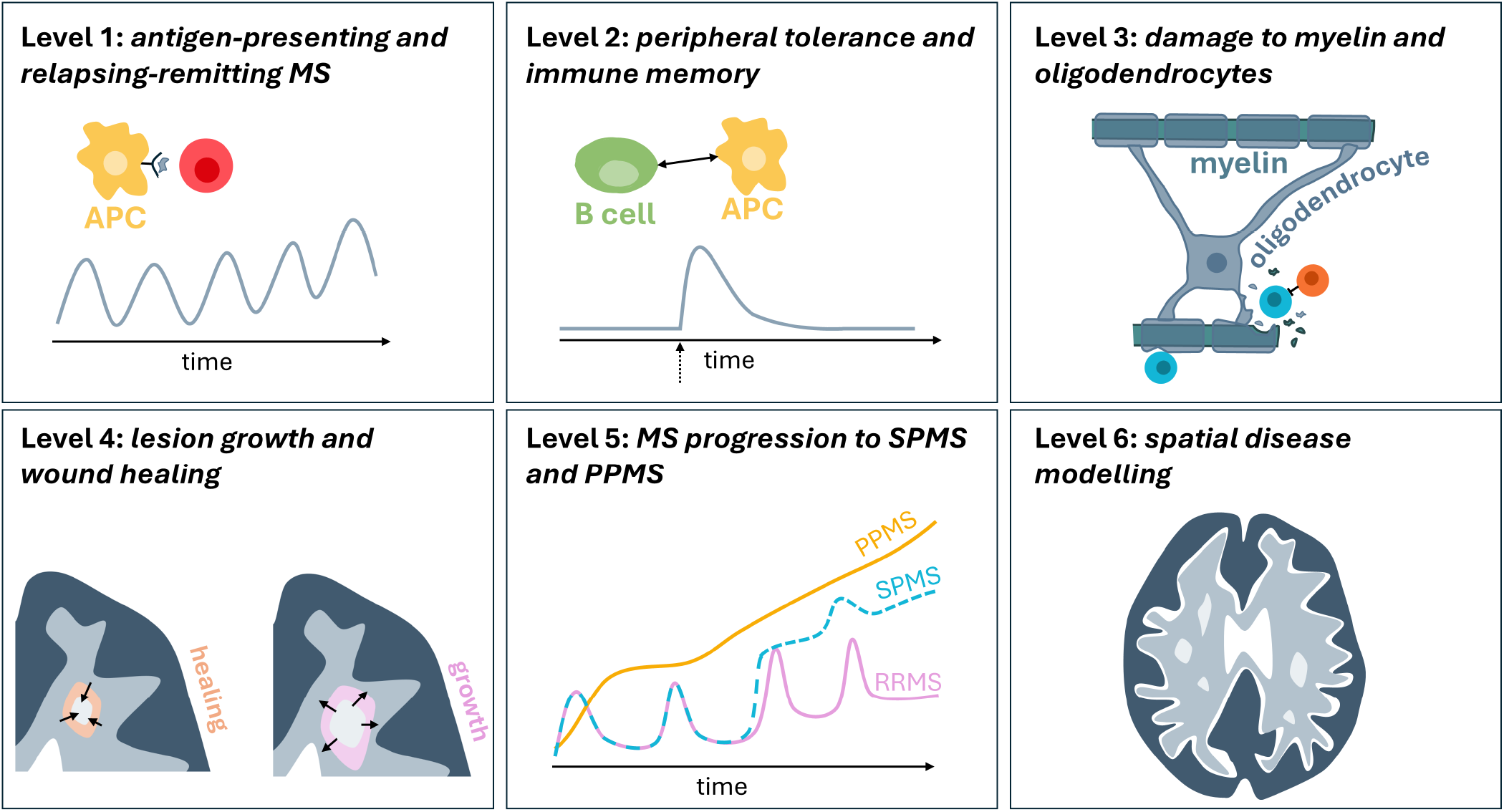

**Highlights:**

- We present a modelling paradigm for MS using systems of ordinary differential equations
- Based on biological principles, we propose six consecutive modelling levels and develop the first three model levels here.
- We investigate the link between risk factors, such as HLA-DR mutations, Epstein-Barr virus infection, estrogen levels, and vitamin D deficiency, to MS expression.

## 1. Introduction

Multiple Sclerosis (MS) is an autoimmune disease that affects the central nervous system (CNS), i.e. brain and spine [1]. Canada has one of the highest MS rates with about 4,000 new patients annually, due to its chronic vitamin D deficiency [2]. MS is characterised by inflammation and neurodegeneration in the CNS [1], which can lead to severe physical or cognitive disability [3]. To date, no cure for MS exists, but medications are available to slow its progression [4]. Clinically, MS manifests itself as visible lesions in the CNS. In these lesions, the myelin coating of the nerve fibers is damaged and scar tissue has formed [5]. Since myelin acts as an electrical insulator of nerve fibers, damage to myelin can lead to significant malfunction of nerve connections [6].

The myelin coating of nerve fibers is an extension of the cell membrane of oligodendrocytes: glial cells in the CNS which help organise brain structure. The loss of myelin in the central nervous system is a normal process, where the loose myelin is removed by microglia, a CNS-resident immune cell. In healthy tissue, oligodendrocytes would fix the missing myelin pieces. In the diseased state, the repair is inhibited due to damage to oligodendrocytes. In fact, oligodendrocyte transplantation is considered a new potential treatment for MS [7]. The damage to oligodendrocytes results from an autoimmune attack [8, 6, 5]. As in many other autoimmune diseases, some self-antigen presenting cells (for example dendritic cells), stimulate the immune response to become active against oligodendrocyte specific cell markers. As the self-stimulation of these self-antigen presenting cells progresses, so does the disease.

There are three main subtypes of MS: relapsing-remitting MS (RRMS), secondary progressive MS (SPMS), and primary progressive MS (PPMS), see Figure 1A. Approximately 85% of patients will initially have RRMS, which is characterised by periods of active disease followed by periods of remission. Around 15-30% of RRMS patients will convert to SPMS [9]: characterised by a worsening of MS disease and steady progression of symptoms. A small group of patients will develop progressive MS, i.e. PPMS, from disease onset, with steady progression of neurological symptoms without periods of remission [10]. A rare, severe type of PPMS is progressive-relapsing MS (PRMS), where individuals experience progressive disease from the onset, as well as flare-ups in disease symptoms [11]. The different phases of MS behaviour (relapses vs progression) are not necessarily distinct, and some patients experience SPMS in their disease, and some patients experience RRMS for their entire disease course.

**Figure 1:**
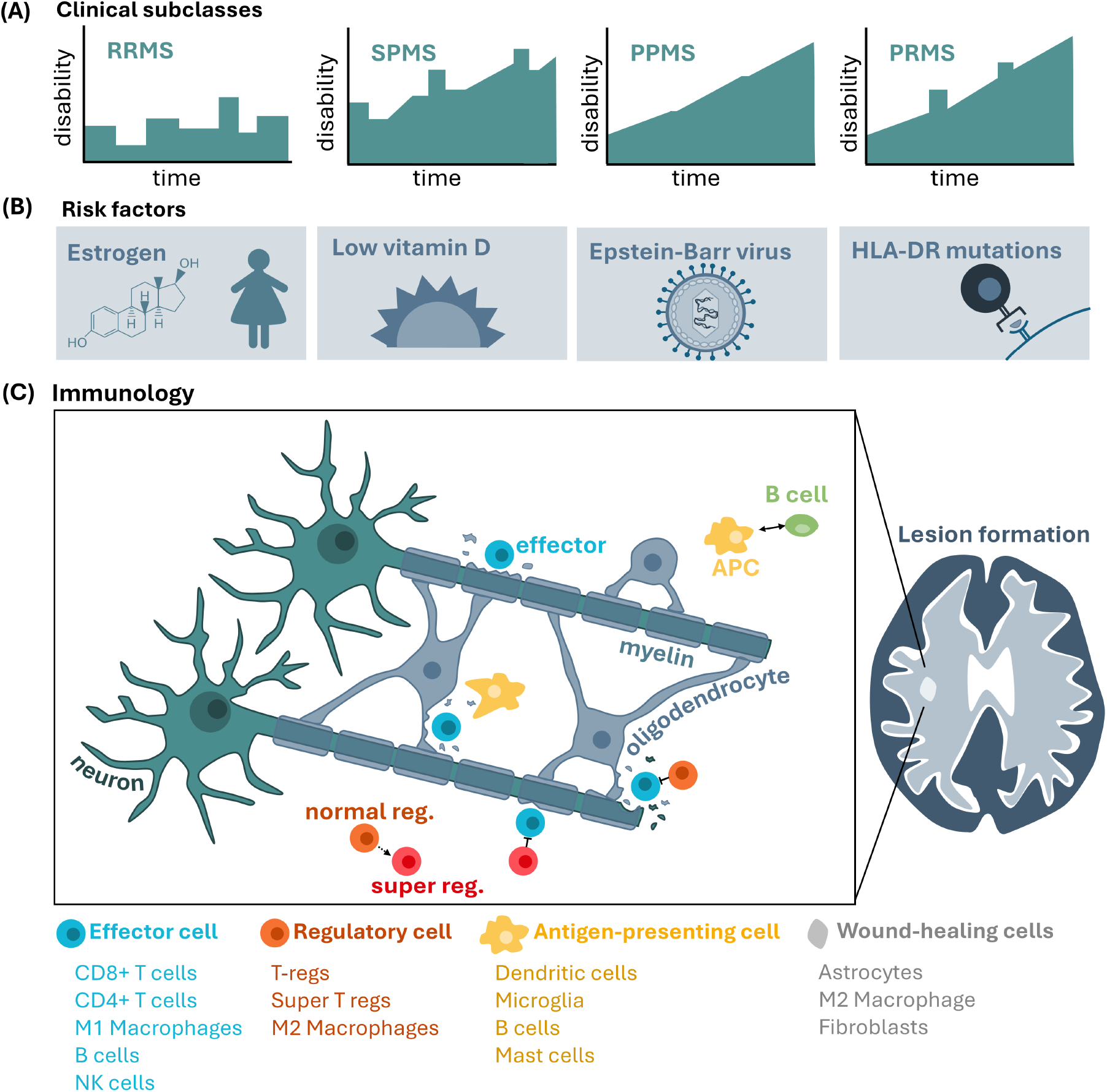
Summary of the MS disease aetiology motivating this study. (A) The three main subtype classifications for MS disease are relapsing-remitting MS (RRMS), secondary progression MS (SPMS), and primary progression MS (PPMS), with a rare subtype of PPMS known as progressive-relapsing MS (PRMS). These disease classifications largely relate to the presentation of an individual patient’s disability over time. RRMS is characterised by periods of relapses and remittance, whereas SPMS is characterised by relapses and periods of progressive disease. PPMS is characterised by progressive increase in disability. Some PPMS patients will experience relapses (or flare-ups) and these are classified as PRMS patients. (B) There are a number of risk factors associated with MS disease and in this work we focus on low/high estrogen, low vitamin D, Epstein-Barr virus (EBV) infection and HLA-DR genetic mutations. (C) The immunology of MS is extremely complicated, in this work we consider a simplified version that incorporates normal and super regulatory cells (regs.), B cells, antigen-presenting cells (APCs) and effector cells (effector). We incorporate into the models their crosstalk and their relationship with neurons, myelin and oligodendrocytes.

Main risk factors associated with MS are Epstein-Barr virus infection, low and high estrogen, and low vitamin D, see Figure 1B. Recent studies have provided compelling evidence for the causal role of EBV infection in MS [12, 13], however, the mechanisms by which EBV contributes to MS pathogenesis are unclear. Vitamin D is considered an environmental risk factor in MS and lifestyle changes and latitude are thought to be driving the increase in MS incidence [14]. It is also well known that there is a sex bias in MS, with a higher incidence in women, which has been linked to fluctuating estrogen [14]. The highest genetic risk factor associated with MS is in the HLA alleles, specifically HLA-DRB1*15:01 [11].

MS is usually diagnosed when typical symptoms are observed, for example walking difficulties, extreme fatigue, numbness or tingling. This can happen at a relative late stage and often the disease will have developed unrecognized for a significant amount of time. This early phase in which disease goes undetected is called the *MS prodrome* [15]. Some early biomarkers for MS prodrome are discussed in the literature such as raised plasma levels of neurofilament light chain (NfL), or the radiologically isolated syndrome (RIS). With RIS, we denote MS-like lesions in brain MRI scans, where the scans were performed for reasons other than MS.

In advanced disease, large areas of demyelination and inflammation are visible in MS patients in the form of lesions observed through MRI [16, 17], see Figure 1C. These lesions are generated through the interaction between oligodendrocytes, myelin, and a number of immune cells: antigen-presenting cells (APCs), B cells, and regulatory and effector cells. The remittance-relapse phase is seen in scans as a series of larger and smaller lesions over time. Also, the geometric structure of lesions suggest a connection between different lesions. This connection is likely facilitated through nerve fibre connections such as white matter fibres in the brain. It is an open question to understand the spatial structure of MS lesions and to substantiate the hypothesis that transport along white matter fibres is essential.

The mathematical modelling of MS is still very young and only a few models are available [18]. From our description above it is clear that several biological scales are involved in the disease progression. The immune systems’ attack on myelin occurs on a microscopic scale, while brain lesions are measured on a macro scale. Here, we present a multiscale framework for understanding MS by focusing on the basic processes of self-antigen presentation, immune stimulation, and immune regulation. We propose a simple ODE-based model that can explain many of the well known effects that relate to MS risk factors such as the influence of Epstein-Barr Virus, the impact of vitamin D deficiency as well as estrogen levels, and the role of the HLA-DR mutation on the disease progression. Although relatively simple, the model does provide new insights into the basic mechanisms at play in MS.

### 1.1. The Multiscale Modelling Paradigm

The literature on the immunopathology of MS is vast, and excellent reviews are available in [8, 6, 19]. Disease progression involves a large number of immune cell types, including macrophages, microglia, T cells, B cells, dendritic cells, and natural killer (NK) cells; as well as CNS-resident cells such as oligodendrocytes, astrocytes and neurons. To complicate matters, one cell type can perform different functions simultaneously. For example, B cells can be self-antigen presenting, they can be memory cells, they can produce antibodies and become effector B cells [20]. In addition, a specific function can be performed by several cell types: for example, dendritic cells, microglia, macrophages, and B cells can all be self-antigen presenting. See Figure 1C for a summary of the main immunology considered in this work.

To write down a mathematical model for all these cell types and all their functions seems impossible. Moise and Friedman [21] published a system of 33 PDEs (partial differential equations) with over 100 model parameters to describe this process. However, it is challenging to parameterise a model with the limited data available. As such, the modelling of MS turns out to be very difficult, and new ideas are needed. Hence here, we take a different approach. Instead of focussing on each specific cell type and their specific cytokine signalling, we take a multiscale approach to identify and model basic processes [22].

Based on the detailed immunological description of MS [8, 21, 5], we propose to divide MS into six subprocesses, each describing a different feature of the disease. Of course, these sub-processes are connected, but we believe that this classification is a useful tool to build a hierarchy of mathematical models for MS, from simple to complex. We show an illustration of these levels in Figure 2.

**Figure 2:**
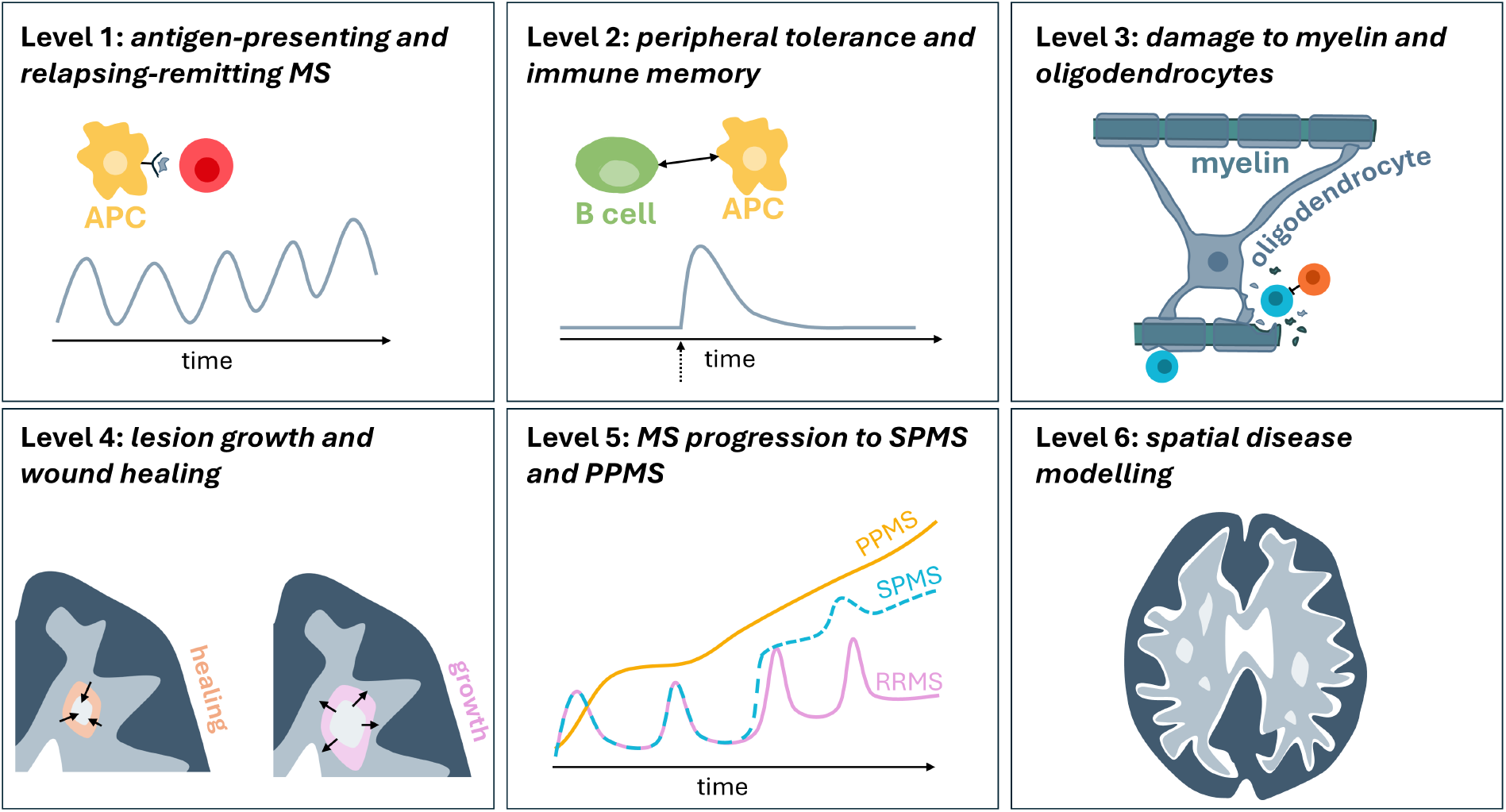
Pictorial summary of Levels 1 - 6 in the modelling paradigm. Level 1 focusses on capturing the basics of the immune response in MS (antigen-presentation and immune regulation) as well as a key characteristic of MS disease: oscillations. Level 2, captures peripheral tolerance (the resilience of the system to small perturbations) and introduces the memory B cell involvement in MS disease. Level 3, incorporates the resulting damage on myelin and oligodendrocytes by the immune response. Level 4, would extend Levels 1-3 to look at how the resulting damage leads to lesions which can grow or shrink over time. Level 5, looks to capture the long-term clinical characteristics of MS: RRMS, SPMS and PPMS. Level 6 would extend Levels 1-5 to capture the spatial component of MS disease and how this influences the immune cell behaviour.

#### MS Level 1: Self-antigen presentation, relapsing-remitting MS and MS prodrome

Level 1 focusses on capturing the autoimmune response based on self-antigen presentation and the resulting oscillatory dynamics of relapsing-remitting MS. A model by Zhang, Wahl and Yu [23] for autoimmune diseases will form our base model here. It balances the activities of antigen presenting cells, effector immune cells and regulatory immune cells. At this level, the effector immune cells are not specified, and are to be understood as a combination of various immune active cells such as macrophages, NK cells, B cell, T cell etc.

#### MS Level 2: Peripheral tolerance and immune memory

The regular overturn of myelin in the CNS is a natural part of brain homeostasis. In the healthy brain, myelin antigens are presented to adaptive immune cells, but no autoimmune response is triggered. This is called *peripheral tolerance*, and a good mathematical model needs to include this case as a healthy control case. As we will see in the analysis that even small initial antigen concentrations in model Level 1 will trigger the full disease (if the parameters are right). Hence the model from Level 1 does not allow for peripheral tolerance. To allow for peripheral tolerance we include memory cells, specifically memory B cells. These are stimulated by myelin related antigens, but they stay dormant if the antigen presentation is low. Only if they are reactivated by a large event will the disease arise.

#### MS Level 3: Damage to myelin and oligodendrocytes

Through activation by antigen-presenting cells, effector and regulatory immune cells damage myelin and oligodendrocytes and this level aims to capture this damage. Upon myelin and oligodendrocyte damage, more self-antigen is released into the environment, allowing antigen presenting cells to show higher concentrations and boosting the immune response further. This likely contributes to the snow-ball effect of a gradually worsening disease and the formation of lesions: areas of inflammation and demyelination visible on MRI scans.

#### MS Level 4: Lesion growth and wound healing

Once the processes of Level 1, 2, and 3 are of significant size and a lesion is established, wound healing mechanisms are engaged [5]. Some macrophages transition from M1 to M2 type, where M2 macrophages focus more on tissue regeneration. Also astrocytes and fibroblasts are activated, which help to regenerate ECM. However, the fine balance of nerve fibres and stroma is destroyed and fibrosis or scar formation might occur, permanently damaging the tissue.

#### MS Level 5: MS progression to SPMS and PPMS

This level focusses on capturing the different long-term clinical stages of MS. All processes 1, 2, 3, 4 together will stimulate a full immune response. Particularly T-cells, for example Th1, Th17 [6], become active, they attack not only oligodendrocytes but also astrocytes, neurons, and macrophages. Moreover, they spread into the periphery of the lesions, invading into healthy brain tissue and causing additional wide spread damage. This stage is likely related to later progressions such as PPMS and SPMS.

#### MS Level 6: Spatial disease modelling

All of the dynamics take place in the spatially heterogeneous environment of the brain and the nervous tissue. Blood vessels, white matter fibres, and gray matter regions all will have different properties for cell movement, cytokine distribution and cell proliferation. To be able to compare model out-puts with MRI scans, it will be necessary to move some of the previous models onto a spatially explicit level.

Mathematical modelling has the advantage that we can develop models on these Levels one by one and successively build up a full MS model. On each level we can then test and see if the model describes some qualitative properties of the disease, and where its limitations are. In this paper, we fully develop models for Level 1 and 2, and start the modelling of Level 3. We test these models on a series of questions outlined below and summarized in Figure 3.

**Figure 3:**
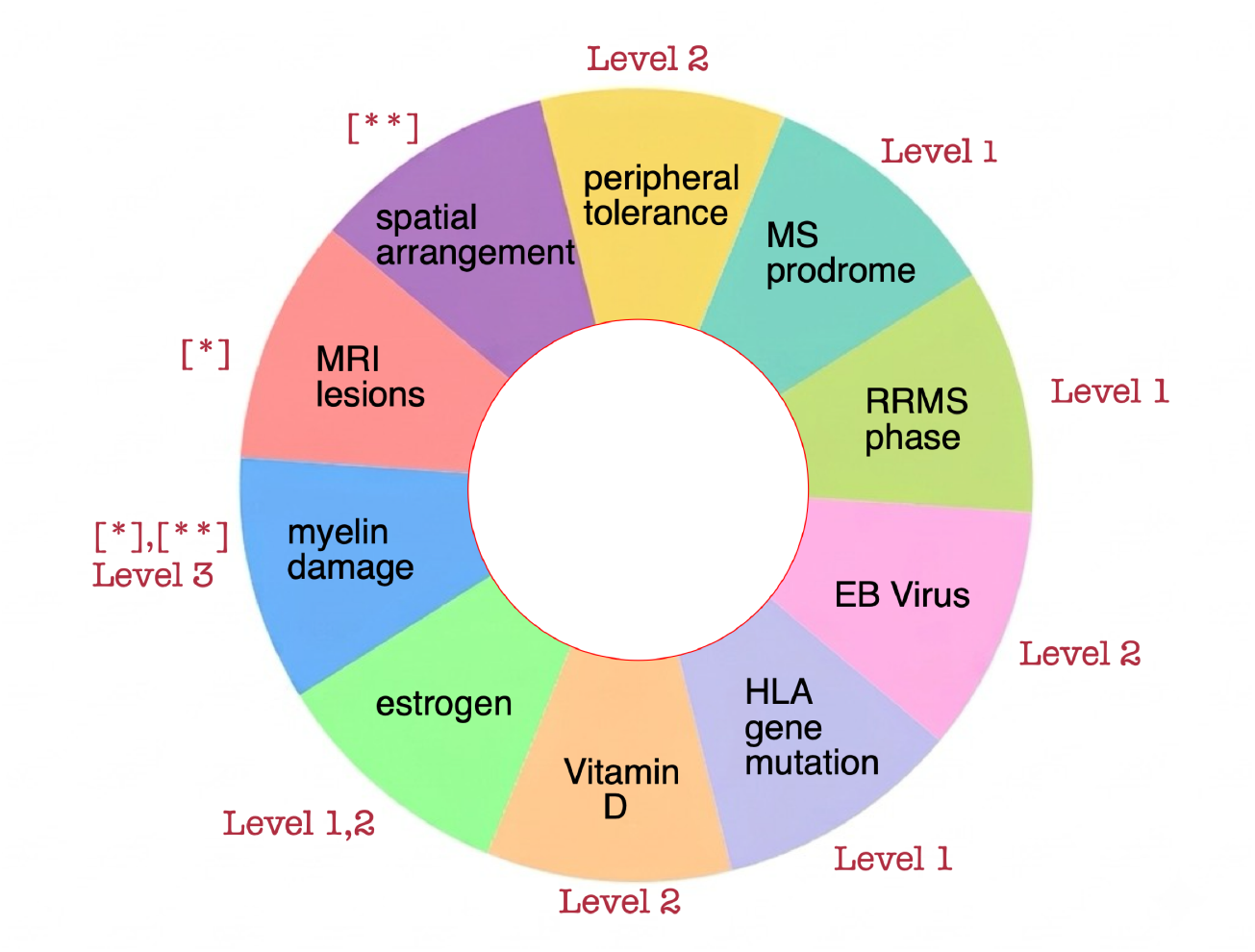
Summary of the models for the biological processes in questions (Q1)-(Q10). [*] = Moise and Friedman [21], [**]= Khonsari, Calvez [25, 26] and more [28, 29, 30, 31, 32, 33]

(Q1) Can the model explain the healthy state of **peripheral tolerance**?

(Q2) Can the model capture the **MS prodrome**?

(Q3) Can the model describe short outbreaks followed by long transients in the **RRMS phase**?

(Q4) Can the model be used to analyse the impact of **Epstein-Barr Virus** [14, 24], or the cytomegalovirus [14], as possible initiator of the disease?

(Q5) Can the model include the effect of mutations, for example in the **HLA-DR gene**?

(Q6) Can the model be used to understand the role of **vitamin D** deficiency [14] on MS?

(Q7) Can the model explain the effect of high and low **estrogen** [14] on MS?

We will see that our simple models of Level 1 and 2 can already provide some insight into these questions. Further questions include,

(Q8) Can the model describe **damages to myelin and oligodendrocytes**?

(Q9) Can the model describe the formation of **MRI-lesions**?

(Q10) Can the model describe the **spatial distribution** of lesions in MRI images?

### 1.2. Previous Modelling of MS

Most previous mathematical modelling of multiple sclerosis has used systems of partial differential equations (PDEs) and focused on Level 6; the spatial appearance of the lesions. Quite influential is the model that was proposed by Khonsari and Calvez [25, 26] in 2007. Their model consists of a coupled system of three partial differential equations (PDEs) for microglia, a signalling chemokine, and damaged oligodendrocytes. The damaged oligodendrocytes release an attractive chemokine and microglia locate at high concentrations of this chemokine via chemotaxis. Using our classification above, this model uses processes from Level 3 and Level 6. Khonsari and Calvez used the model to describe Balo’s MS, which is a rare form of MS that shows concentric rings of demyelination patterns [27]. Their model was able to describe these patterns [25, 26].

The model of Khonsari and Calvez [25, 26] has subsequently been extended and analysed in a series of papers, most notably those from the Lombardo group [28, 29, 30, 31, 32, 33]. The focus of these papers lies on a full mathematical analysis of Khonsari and Calvez’s model and extensions, including results on local and global existence, on spatial pattern formation, and underlying bifurcation analyses. The model of da Paula *et al*. [34] further extends the model to focus on two compartments, the brain parenchyma and the lymph nodes. In their model they consider a spatial model in the brain region and an ODE model in the lymph nodes. The lymph node model is used to describe self-antigen presentation to the immune response and the subsequent immune activation. The latest addition to this line of work is a new paper from Bisi *et al*. [35] where classes for self-antigen presenting cells and regulatory immune cells are added to the Khonsari-Calvez model. Again, the focus lies on spatial pattern formation, and a large variety of spatial patterns are found, including stripes, rings, and hexagonal patterns.

Another important contribution is the model of Moise and Friedman [21] mentioned earlier. They developed a detailed model for MS including all relevant immune cells and most of the known cytokines, resulting in a system of 33 coupled PDEs. Their model includes a free boundary, which describes the spread of lesions as driven by myelin and oligodendrocyte damage. Compared to our classification above, this model encompasses all Levels from 3 to 6. In this work, however, we propose that a model for Levels 1 to 6 can be built with a reduced level of complexity.

The work mentioned above all use differential equation models. There are many other approaches for mathematical modelling of MS, such as stochastic models, statistical models, and agent based models [36, 37, 38]. Several of these have been developed for MS and the comprehensive review article of Weatherley *et al*. [18] provides an extensive list of references. Recent work by Kennedy *et al*. [39] used patient data to inform a multilayer network model of MS and identify connections between molecular aspects of MS and overall phenotype.

We summarize the main results of our modelling, and some previous models, in relation to the questions proposed in this paradigm in Figure 3.

## 2. MS Level 1: Self-antigen presentation, MS prodrome, and RRMS

### 2.1. Level 1 model

A key feature of the initial stage of MS is the relapse-remittance cycle (Figure 1A), which can persist for many years. Researchers have attempted to describe these cycles through various models that are based on a predator-prey interactions. For example, Eletterby and Ahmed [40] use a predator-prey type model where oligodendrocytes are prey and immune cells are the predator. The model is, in fact, very similar to standard models for oncolytic virotherapy [41], and it is well known that these models show a Hopf bifurcation for appropriate parameter values. However, this predator-prey dynamic in MS assumes a high self-renewal rate of oligodendrocytes, when in reality the self-renewal of oligodendrocytes occurs very slowly at 0.3% per annum in a healthy individual [42].

A model by Zhang, Wahl and Yu [23], which will be the basis of our model, describes the cyclic phase of MS and other autoimmune diseases through the interplay of self-antigen presenting cells (such as dendritic cells, or microglia), effector immune cells and regulatory immune cells. Their model considers the activation of effector cells through self-antigen presentation, the subsequent immune response, and the suppressive action of the regulatory immune cells. In particular, Zhang *et al*. [23] recognize the heterogeneity of the regulatory immune cells, which show sub-populations that can be distinguished by their expression of the HLA-DR receptor. For example HLA-DR^−^ T cells are immature, long lived and have a reduced effect. HLA-DR^+^ T cells however, are terminally differentiated, highly effective at controlling T cells, and short-lived. Similarly, macrophages appear in a range of phenotypes of which the M1 type macrophages are more inflammatory while the M2-type macrophages are more regulatory. Again, we do not attempt to focus on one specific class of regulatory cells, but we consider them as a regulatory cell compartment where we allow for two regulatory mechanisms (as done in Zhang *et al*.) and call them *T* (*t*) for “normal” regulatory cells and *S*(*t*) for the stronger regulatory type (or “super regulatory cells”).

The original model of Zhang *et al*. [23] is an ODE of five equations for the self-antigen presenting cells, one type of effector cells, and two types of regulatory immune cells, and the antigen. They show that the model has a Hopf bifurcation and oscillating solutions, which are consistent with the RRMS phase. In addition, they identify fast and slow time scales which invites a multiscale analysis. To leading order, the model reduces to three equations and we use these as our Level 1 model to describe self-antigen presenting cells *D*(*t*), normal regulatory cells *T* (*t*), and stronger regulatory cells *S*(*t*). In non-dimensional form the model reads

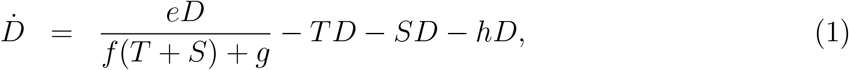

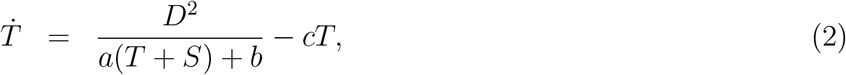

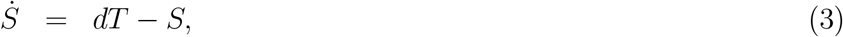

where *a, b, c, d, e, f, g, h* are positive constants. The corresponding flow diagram is shown in Figure 4 and parameter values are in Table 1.

**Table 1:**
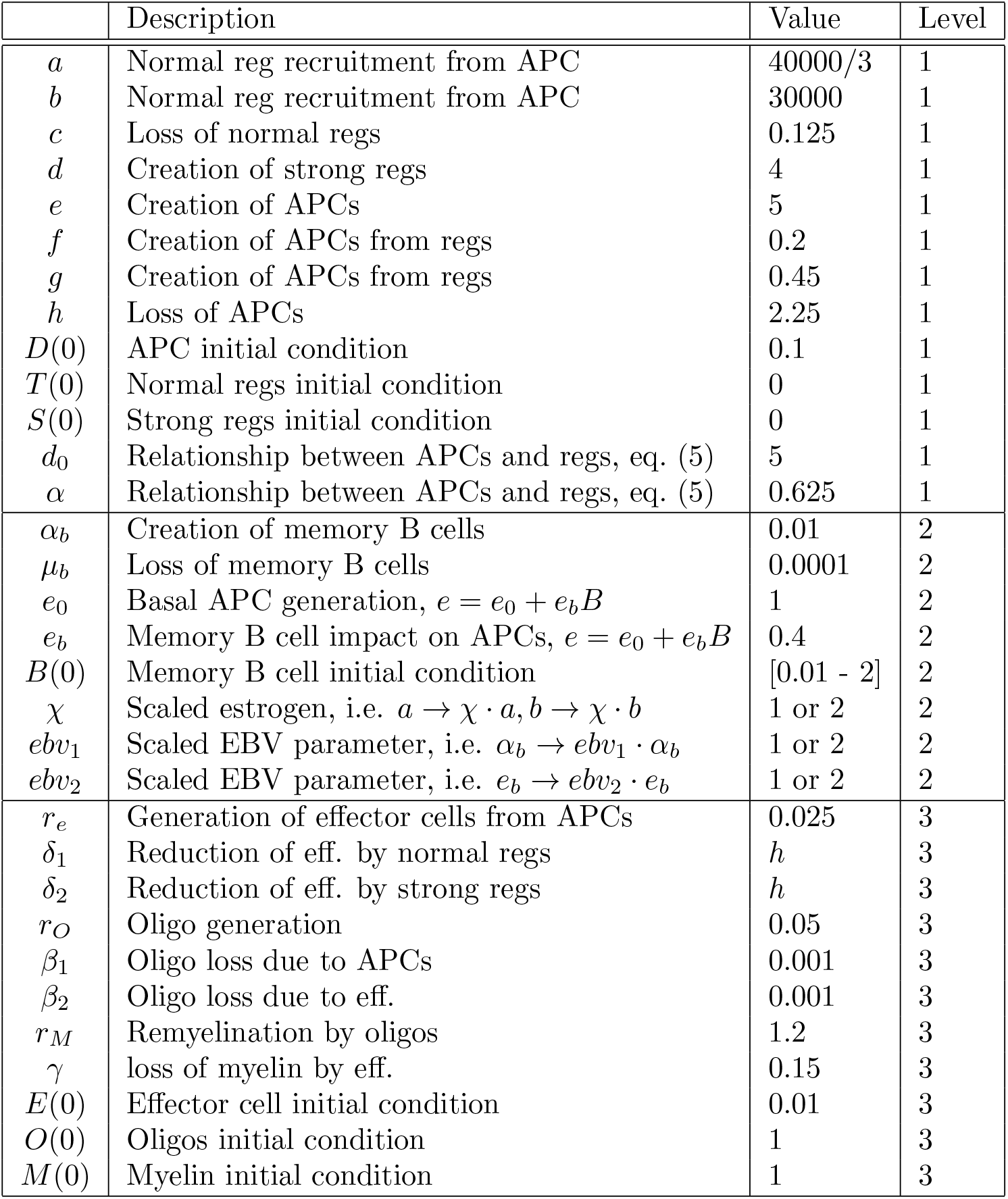
Base parameter values and their meaning for model Level 1, 2 and 3. The parameters for the Level 1 model are taken from [23]. Since the model is already non-dimensionalised, all parameters are dimensionless. The parameters for models on Level 2 and 3 were chosen relative to those of Level 1 to obtain biologically realistic ranges. We describe the parameter choices in detail Section 5.

**Figure 4:**
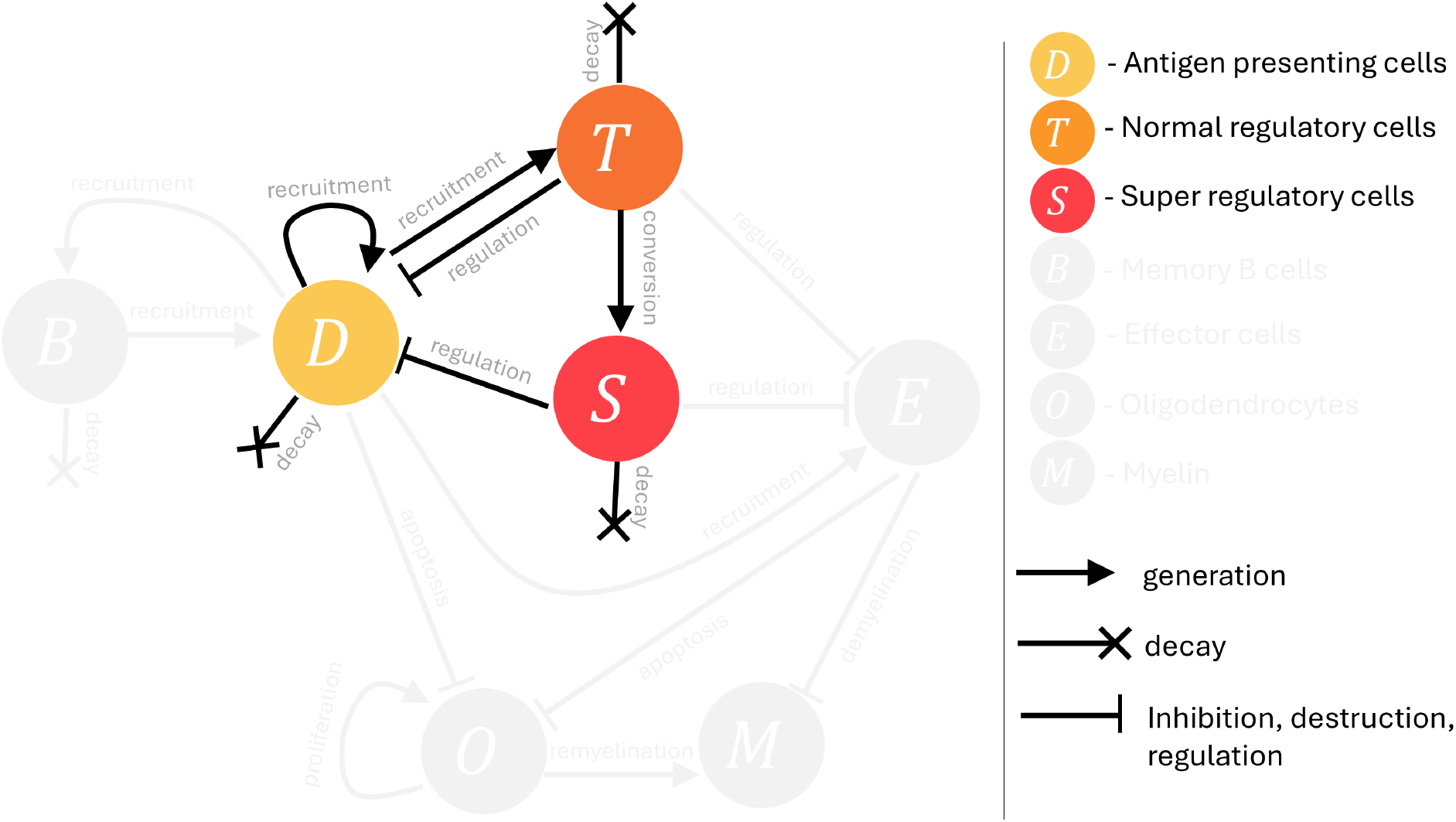
Level 1 model flow diagram for Equations (1)-(3). At this level, the focus is on modelling disease initiation and relapsing remitting dynamics. In this model, antigen-presenting cells (APCs), *D*, are undergoing recruitment following some unspecified catalyst event. APCs recruit normal regulatory cells *T*, which in turn can convert to super regulatory cells *S*. These regulatory cell populations both inhibit APC recruitment. This creates a negative feedback loop between the APCs and regulatory cells. All cells in the system naturally decay. The arrows represent promotion and flat bars represent inhibition, where crosses represent decay. The greyed out section of the model will be introduced in Levels 2 and 3.

The above model in Equations (1)-(3) arose from a scaling limit and a non-dimensionalisation of a full antigen-immune response model in [23]. All variables and rates in Equations (1)-(3) are unitless. If rescaled to dimensional variables, then a time unit corresponds to 5 days. The parameter values from [23] are summarized in the first box of Table 1 and the parameter choices are discussed in Section 5. The self antigen activation rate *e* will be our main bifurcation parameter.

### 2.2. Simulations and analysis of MS Level 1 model

To illustrate the dynamical behaviour of the Level 1 model (Equations (1)-(3) and Figure 4), we perform numerical simulations for the base parameters in Table 1. In Figure 5A, we use a small antigen presentation by choosing a small value for *e* = 1. In that case the antigen presenting cells are not able to generate a significant immune response, and the disease does not start. This healthy state is the normal behaviour for people that do not suffer from MS. In Figure 5B, we use the base value for *e* = 5, which triggers a cycle of autoimmune responses. In Figure 5C we slowly increase the antigen presentation according to the linear model

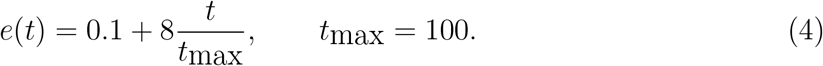

**Figure 5:**
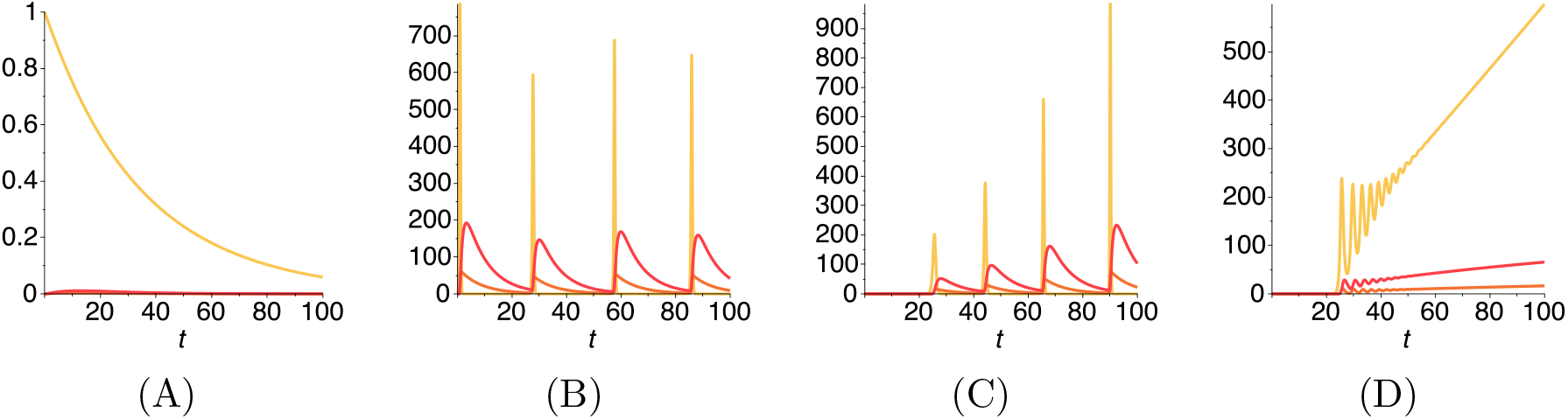
Typical time progressions of the Level 1 model, Equations (1)-(3), for the parameters in Table 1. The colours are yellow for *D*, orange for *T* and red for *S*. (A) Healthy state for low antigen presentation *e* = 1.i.e. no MS disease. (B) The first disease appearance for *e* = 5: high enough antigen presentation triggers a remittance-relapse cycle. (C) Onset of disease after 25 time units where *e*(*t*). (D) Increase in antigen presentation and exhaustion of regulatory T cells where *c*(*t*). Note in (A) the curves for *T* and *S* were multiplied by 100, to make them visible. In (B), (C), (D) they were multiplied by 20.

This case mimics a scenario where antigen presentation builds up over time. This mechanism could be a proxy for the major histocompatibility complex (MHC) Class II genes, particularly those related to HLA-DR (human leukocyte antigen DR isotype) that dominate genetic susceptibility in MS [43, 44, 45]. Genetic factors related to this gene are believed to influence the way peptide antigens are presented to CD4+ T cells for immune recognition. [14]. In addition, these alleles shape a CD4+ T cell repertoire that is more strongly responsive to MS-associated infectious organisms and autoantigens [46]. Another possible mechanism for this form of *e*(*t*) are memory B cells [20], which we will discuss in the next section for the Level 2 model. After a delay of 25 time units, the RRMS cycle starts with increasing intensity.

Finally, in Figure 5D, we increase the antigen presentation, and at the same time, we include exhaustion of the regulatory T cells. We do this by increasing the death rate of the *T* cells as

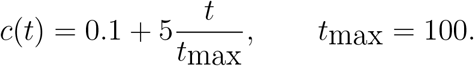

In this case the disease starts again at time unit 25, progresses through a few cycles until it starts growing uncontrollably. This shows a scenario that might explain the transition from RRMS to PPMS and SPMS; however, we leave exploration of this to later model levels.

### 2.3. Analysis of the model test questions (Q1)-(Q9)

We use the Level 1 model in Equations (1)-(3) to test our research questions (Q3), (Q5), and (Q7).

#### (Q3)

The most striking property of Equations (1)-(3) is the robust expression of periodic outbursts once the bifurcation parameter *e* is large enough. The lengths of the gaps between outbursts depend on the parameters, and long periods of quiescence are possible. This property forms the back-bone of our further modelling extensions.

#### (Q7)

High levels of estrogen sensitize the immune system to any type of invader [47, 48]. This means the immune system is also more sensitive to self-antigen presentation such as myelin or oligodendrocyte related antigens. In our model, this would correspond to an increase in the antigen presentation parameter *e*. We saw in our simulation in Figure 5A and B that an increase in *e* can lead to disease outbreak. Of course, female estrogen levels are not permanently high, as they go through the menstrual cycle. Still, there are phases with increased risk.

#### (Q5)

Mutation in the HLA-DR receptor gene can lead to enhanced sensitivity to self-antigen presentation [46]. This again affects the parameter *e* in our model. As mutations might accumulate over time, we can use the case of increasing *e*(*t*) from (4) as an illustrative example. As seen in Figure 5C, the disease outbreak can be promoted by this mutation.

### 2.4. Bifurcation Analysis of Model Level 1

We have seen in the previous simulations, and also in the work of Zhang-Wahl-Yu [23], that model (1)-(3) has a Hopf bifurcation and stable oscillations. Any of the model parameters can be used as a bifurcation parameter, and Zhang-Wahl-Yu used *e, c* and *d*. There is no need to repeat it here, instead, we consider a biologically meaningful combination of parameters *d* and *e* to illustrate the generation of two Hopf bifurcations.

We assumed that super regulatory cell generation from normal regulatory cells would depend on the strength of APC recruitment, i.e., as APC recruitment increases, we would see a decrease in the creation of super regulatory cells: *d* describes the creation of super regulatory cells, *S*, from normal regulatory cells, *T*, and *e* describes the APC recruitment. We fixed a linear relationship between *d* and *e* such that as the APC recruitment increased, we’d see a decrease in the creation of new super regulatory cells, i.e.,

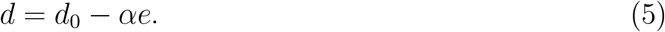

Biologically, this suggests that when the APC recruitment approaches zero, i.e. *e* → 0, we have a maximum conversion, *d*_0_, of regulatory cells from normal to super (their terminal counterpart). This simulates the dampening down, or slowing down of the inflammatory response, and transition to a regulatory immune response. Conversely, we assume that as the recruitment of APCs is increasing *e* ≫ 0, we see a slowing of the transition of normal to super regulatory cells, suggesting a prioritisation of the normal regulatory population. Specifically, APCs activate effector T cells and regulatory T cells (Tregs) are then activated and recruited to control the effector functions [49].

Fixing the linear relationship between *d* and *e* in Equation (5), we then examine the impact of increasing the APC recruitment slowly over time. In other words, we conduct a one-parameter bifurcation analysis for the parameter *e* using Equations (1)-(3), see Figure 6A-C. We observe a Hopf-bubble as *e* increases: first a Hopf bifurcation gives rise to stable limit cycles, these eventually die away with a second Hopf bifurcation. Conducting a two-parameter bifurcation analysis we can see the existence of a region of limit cycles in (*e, b*), (*e, c*) and (*e, α*) parameter space, Figure 6D-F.

**Figure 6:**
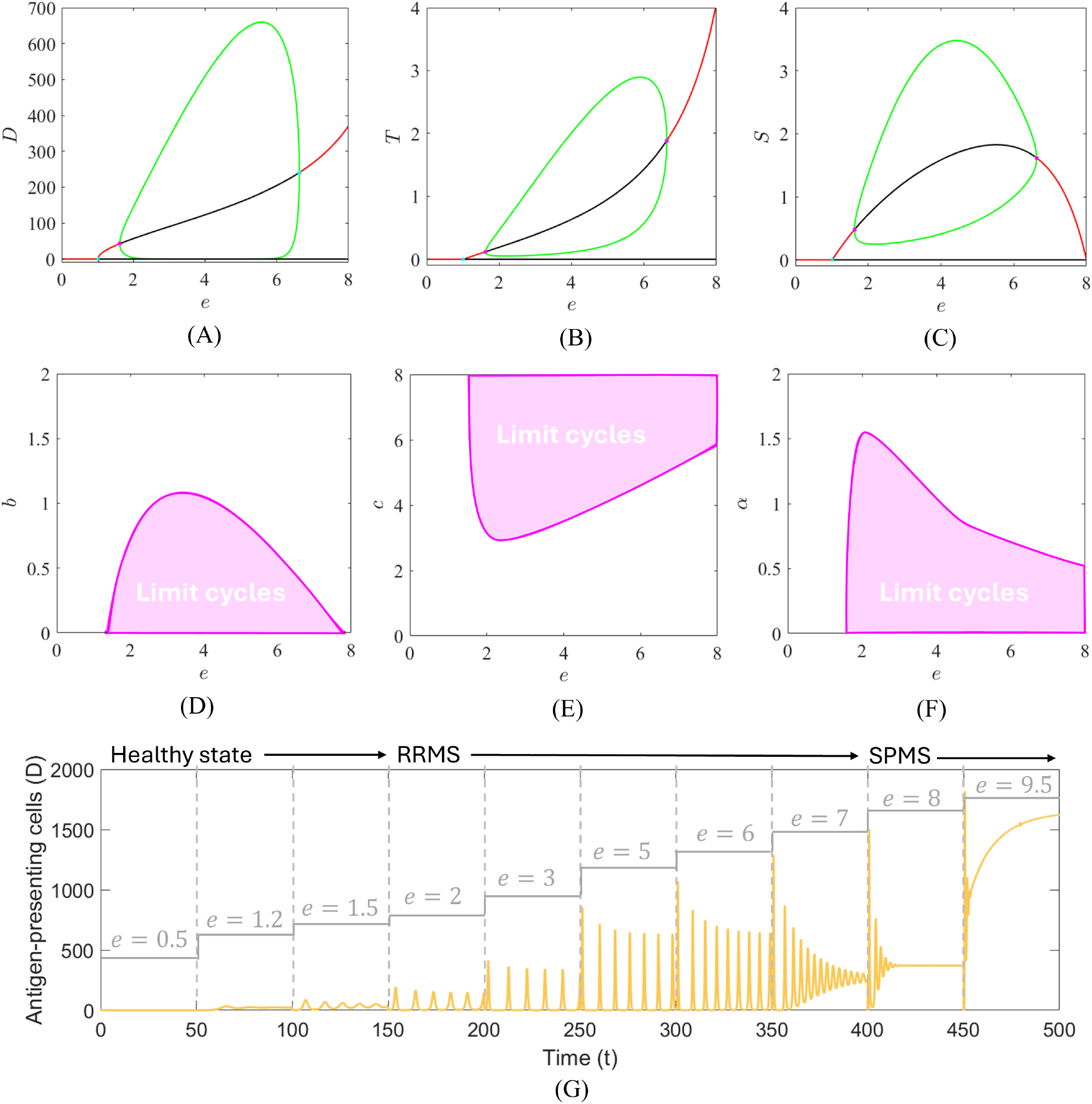
Bifurcation diagram for Equations (1)-(3) with the constraint in Equation (5). (A)-(C) One-parameter bifurcation diagrams for *e* and variable (A) *D*, (B) *T* and (C) *S*. Solid red curves represent a stable steady state and the black curve represents an unstable steady state. Green curves represent the maximum and minimum value of the limit cycle for the variable on the vertical axis. In (D)-(F) are the two parameter bifurcation diagrams for *e* and (D) *b*, (E) *c* and (F) *α*. All bifurcation analysis was performed in XPPAUT and plots were generated using XPPLORE [50]. In (G) We plot *D* over time, varying *e* every 50 time units. Parameters are fixed at *f* = 0.2, *d*_0_ = 5, and *α* = 0.625, and the remaining values are given in Table 1.

To understand the relevance of the appearance and then disappearance of limit cycles for MS disease, we have simulated the model over a longer time period for increasing values of *e*, see Figure 6G. At first, for low values of *e*, we have the antigen-presenting cells returning to zero, and the system exhibiting a healthy stage. At a small level of *e* the system undergoes a transcritical bifurcation and a non-trivial coexistence steady state arises, which is stable for small *e*. We claim that this stable coexistence steady state can be associated with the MS prodrome. It is a state that shows already an increased self-antigen presentation but it has not developed into a full disease [15]. As *e* increases further, we see the transition to relapsing disease, i.e. RRMS. Continuing the increase in *e*, we eventually see the progression from oscillating, relapsing, disease, to a steady increase in antigen-presenting cells, reminiscent of SPMS. This suggests a possible explanation for the evolution of MS disease from a healthy state to RRMS and then SPMS could be through an increasing creation of antigen-presenting cells (*e*) and at the same time, a decrease in super regulatory cells, i.e. *d*. Based on this analysis we can add

#### (Q2): MS Prodrome

The bifurcation analysis of Zhang-Wahl-Yu [23] and our own analysis all show an initial transcritical bifurcation for a small bifurcation value, where a stable coexistence steady state arises. This necessitates the existence of a pre-disease state, which we can identify as the MS prodrome. In this respect, the mathematical model acts as evidence that such an initial disease state must exist.

From the Level 1 model, we have captured the onset and relapsing-remitting dynamics of MS by solely focusing on the interaction between APCs and regulatory cells. Increasing and decreasing the antigen presentation, we have captured the transition from healthy state to MS prodrome, to relapsing-remitting and then sustained disease. However, as soon as the model parameters are in the oscillatory range, a small initial self-antigen signal will be amplified and lead to a full disease response. This is not seen in reality since people with reduced Vitamin D, for example, do not always get MS. The Level 1 model does not account for peripheral tolerance, the effect that a small antigen signal is tolerated and does not lead to a disease. In our Level 2 model, we address this shortcoming by the inclusion of memory B cells.

## 3. MS Level 2: Peripheral tolerance and immune memory B cells

Although MS has long been considered a T cell driven autoimmune disease, evidence to support the involvement of B cells is growing [51]. Interest in B cell phenotypes arose following the efficacy of B-cell depleting treatments of MS patients [20]. Treatments for RRMS patients, such as cladribine and alemtuzumab, have been shown to markedly deplete memory B cells and as a result improve patient symptoms [52, 51].

Memory B cells build up over time once self-antigens are presented. They persist for a very long time and react quickly upon further antigen presentations. Memory B cells can clonally reproduce and can also proliferate into effector B cells [20]. These effector B cells then attack myelin and oligodendrocytes, progressing MS disease and consequently producing more antigen.

### 3.1. Level 2 model

To capture memory B cells in the Level 1 model, Equations (1)-(3), we introduce a new population *B*(*t*). We change the antigen activation rate *e* to *e*_0_ + *e*_*b*_*B*(*t*), with a base rate *e*_0_ and a B cell related activation *e*_*b*_*B*(*t*), and introduce a new equation for *B*(*t*). Memory B-cells are produced through increased self-antigen presentation and then remain for a long time after production. We assume the production is proportional to the rate of increase in *D*(*t*) for 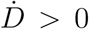. However, when the antigen presentation decreases, 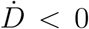, we have no decrease in the B cells (except natural decay). Hence, we use a term max 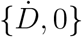 as a growth rate. We sketch the Level 2 model diagram in Figure 7 and our Level 2 model becomes:

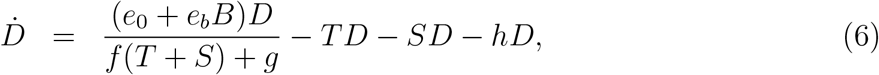

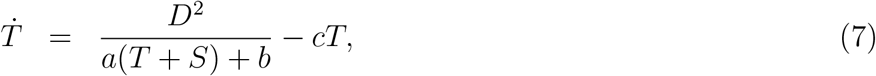

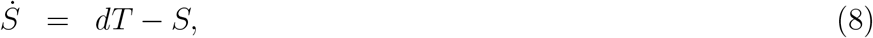

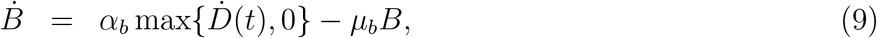

where *α*_*b*_ is a constant of proportionality between the growth of *D* and *B*, and *µ*_*b*_ is a very small natural decay rate for *B*.

**Figure 7:**
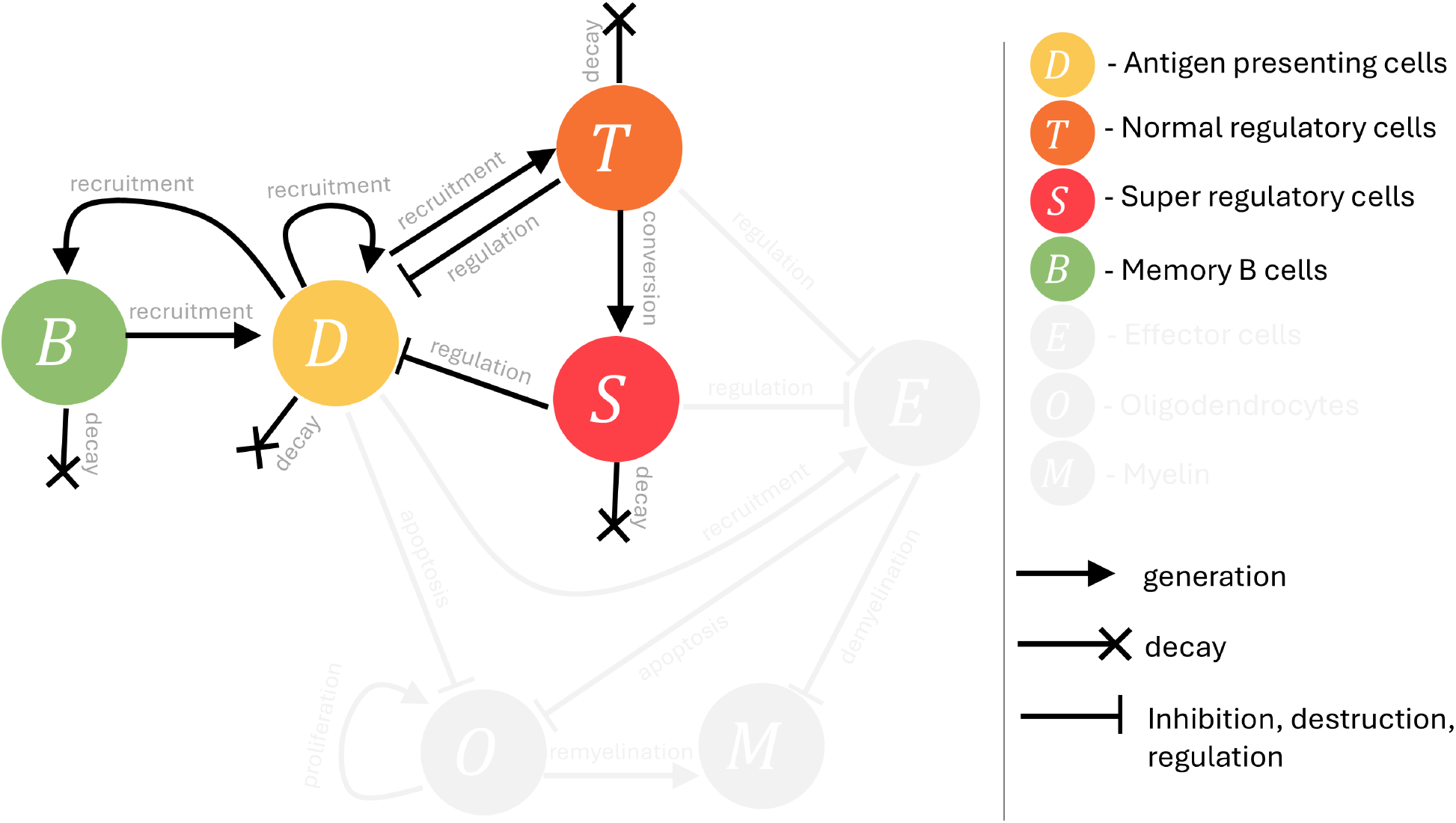
Level 2 model flow diagram for Equations (6)-(9). At this level, the focus is on incorporating memory B cells. In this model, memory B cells *B* have been added to the previous Level 1 model (see Figure 4). Memory B cells are recruited by APCs, and also influence APC recruitment. Memory B cells also decay. Arrows represent promotion and flat bars represent inhibition, where crosses represent decay. The greyed out section of the model will be introduced in Level 3.

### 3.2. Analysis of the model test questions (Q1) - (Q7)

We use the Level 2 model in Equations (6)-(9) to test our main research questions (Q1)-(Q7). The base parameter values are given in Table 1.

#### (Q1) and (Q3): peripheral tolerance and RRMS

To test for peripheral tolerance, we simulate the model for increasing initial condition of memory B cells. The results are shown in Figure 8. For low initial memory B cells *B*(0), the antigen signal (yellow curve) declines and also the memory B cell population (green curve) declines. If *B*(0) is increased from 0.001 to 0.1, we enter a small-level coexistence steady state. The antigen signal does not fully disappear, but it also does not explode in a large scale signal. In the same way, the memory B cells attain a small but non-zero level. Once *B*(0) exceeds 1, we get a full disease response. The antigen signal (yellow curve) enters a periodic response with large outbreaks in regular intervals. At each outbreak, the memory B cell population grows significantly, making each following antigen presentation even more dramatic. In these simulations, we clearly see that the initial amount of memory B cells is the driving force for MS disease progression to larger and larger inflammatory events, while the dynamics of the Level 1 model for antigen and immune regulatory cells explains the oscillations of the RRMS phase.

**Figure 8:**
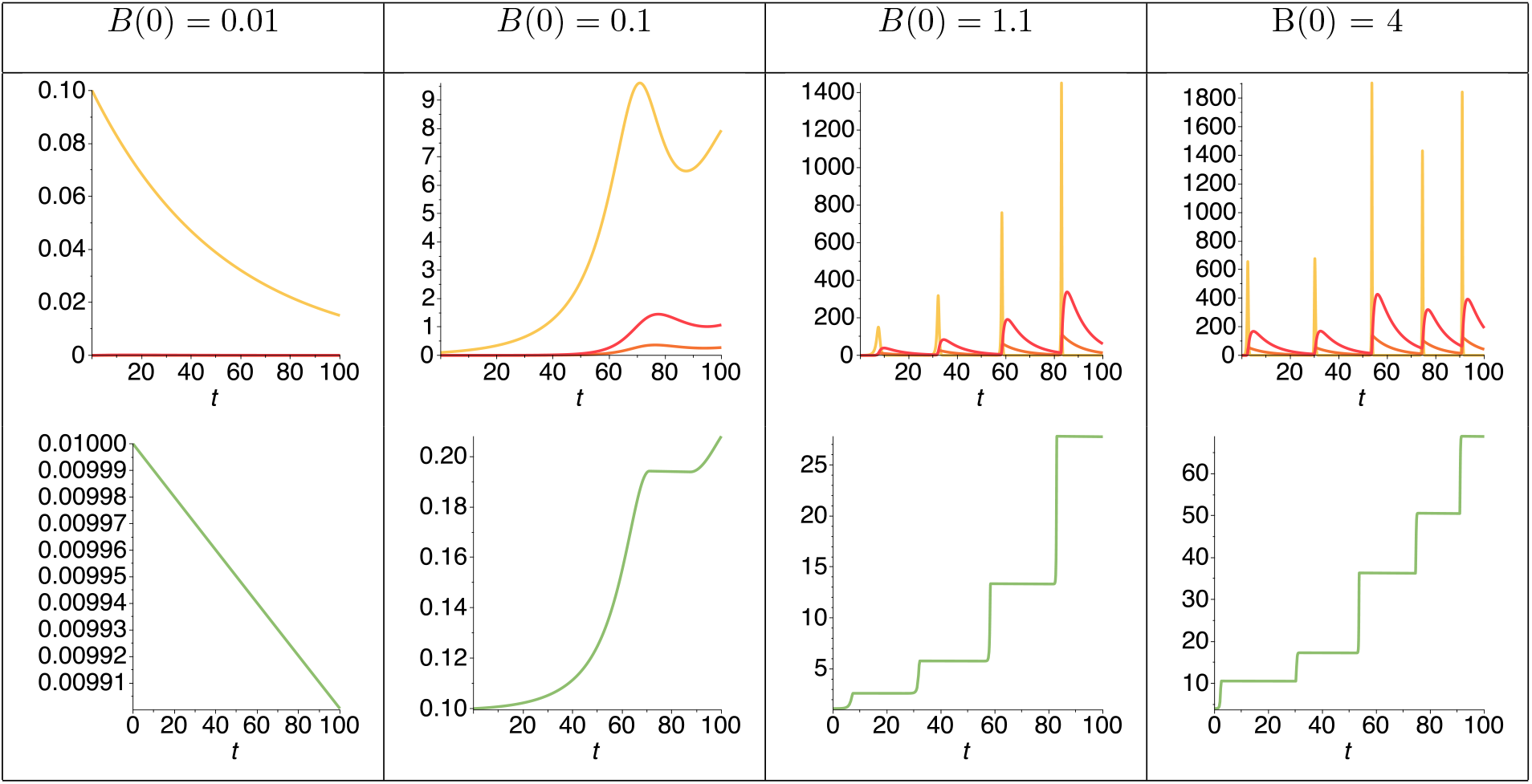
Simulations of Level 2 in Equations (6)-(9) with the parameters identified in the text, for different initial conditions for *B*(0). Top row: *D*(*t*) (yellow), *T* (*t*) (orange) *S*(*t*) (red), Bottom row: *B*(*t*) (green). For small values of memory cells, an initial antigen signal is suppressed and no disease starts, while for larger initial values *B*(0) *>* 1 we obtain a full RRMS response. Also note the continued increase of memory B cells over time in the diseased cases.

The healthy case (peripheral tolerance) corresponds to low values of antigen activated memory B cells. Small events will not amplify and the system will reset to a healthy state. However, if the memory B cell population is significantly enlarged, for example due to a viral infection or some form of trauma, then an irreversible disease response could be triggered.

#### (Q4) Epstein-Barr virus

To include the effects of the Epstein-Barr virus and other MS-related factors, such as estrogen and vitamin D, the parameters *e*_0_, *e*_*b*_, *a, b* will be modified. To model the effect of the Epstein-Barr virus (or cytomegalovirus), we consider two effects [14]. On the one hand, the viral infection directly contributes to the production of memory B cells, hence we consider a factor *ebv*_1_ and use

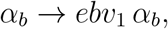

where we consider *ebv*_1_ = 1 or 2.

Secondly, the EBV antigen has been identified to be very similar to myelin related antigens [6, 14] and memory B cells that were activated through the viral infection also react to myelin related signalling. Such an event can increase the initial condition for memory B-cells in our model and it adds to the effect of the memory B cells overall. In our model, we introduce a factor *ebv*_2_ and consider

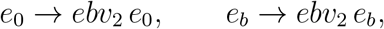

and we consider *ebv*_2_ = 1 or 2.

The results are shown in Figure 9. The first column shows the base case, where no disease response arises and the memory B cells attain a small non-zero homeostasis level. In the second column, we increase *ebv*_1_ from 1 to 2 and see a delayed response after about 100 time units. Indicating that a viral infection could trigger MS disease years after the initial viral infection. In column 3, we increase *ebv*_2_ from 1 to 2, and the response is even more dramatic with a strong disease response and increasing memory B cells.

**Figure 9:**
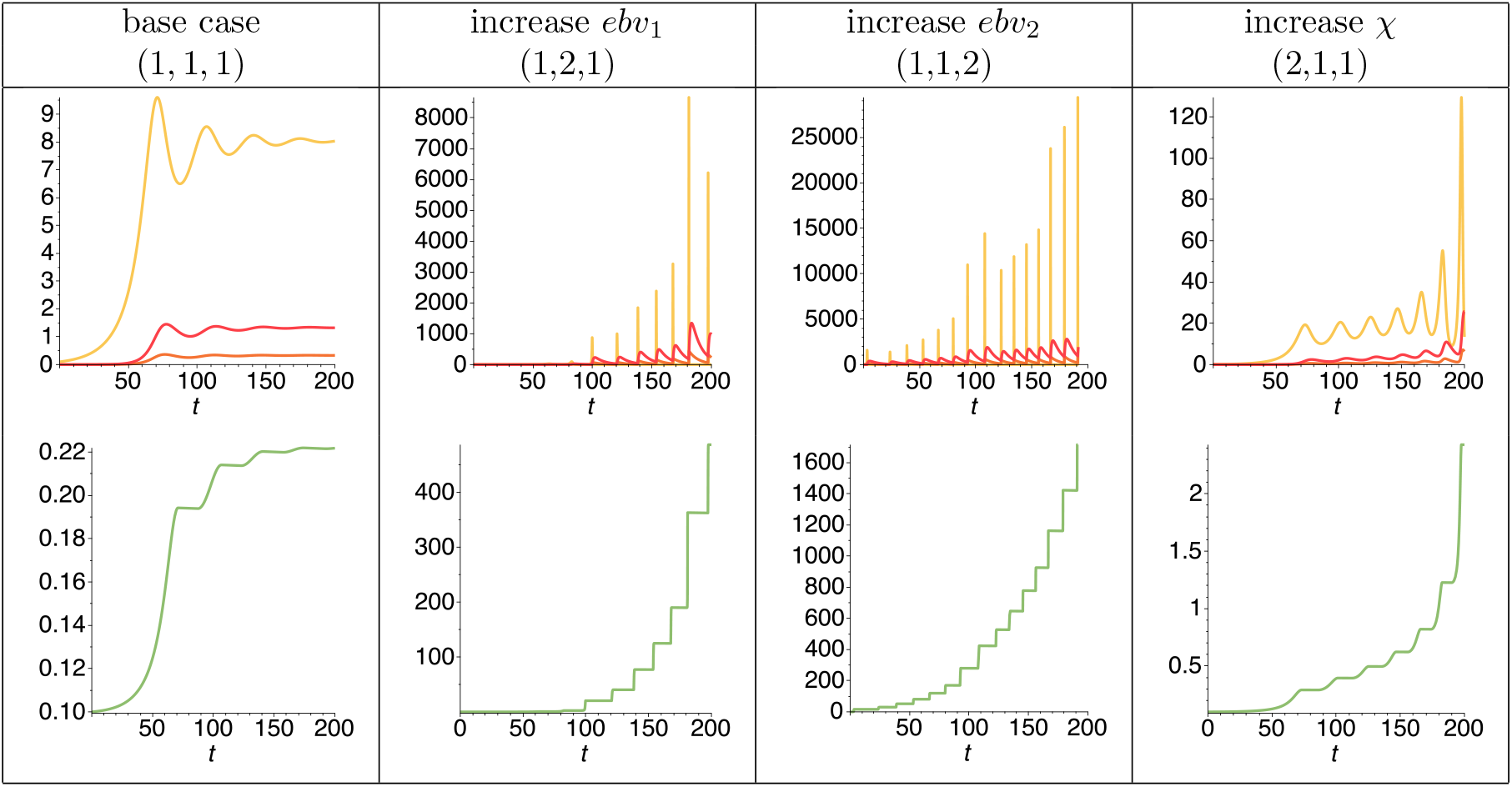
Simulations of Model Level 2 in Equations (6)-(9) as functions of time for *D* (yellow), *T* (orange), *S* (red) in the top row and *B* (green) in the bottom row. The tuple indicates the three parameters (*χ, ebv*_1_, *ebv*_2_). Top row show the cell densities and bottom row the memory B-cells response. Note the different scales on the *y*-axis.

#### (Q6) and (Q7), estrogen and vitamin D

Low Estrogen and low vitamin D have similar effects on the immune response. Vitamin D deficiency and low estrogen down-regulate the regulatory immune response [14, 48]. Hence, we modify the production term of the regulatory cells by a factor *χ* as

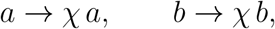

where 1*/χ* describes the fraction of vitamin D or estrogen as compared to the base model. We consider *χ* = 1 and *χ* = 2. The result is shown in the last column of Figure 9. Continued deficiency of vitamin D and estrogen can lead to a long duration of intermediate oscillations, which eventually lead to a disease response.

#### Combined Effects

In reality, the effects that we study here do not act in isolation, but they might combine and amplify each other. Here we consider combinations of two of our model parameters and present their response in heat map diagrams.

In Figure 10A, we vary the initial condition for memory B cells *B*(0) on the *x*-axis and the initial value of the antigen presenting cells *D*(0) on the *y*-axis. Dark red indicates no disease response, corresponding to the peripheral tolerance case. We assumed that the key markers of MS disease would be the presence of more than three oscillations with increasing amplitude and also high levels of APCs. We note that there is a large range of initial B cells and APCs that will give rise to MS disease, suggesting that the system is very sensitive to these initial conditions. In Figure 10B we vary the Epstein-Barr virus parameter *ebv*_1_ on the *x*-axis and the vitamin D and estrogen deficiency parameter *χ* in the *y*-axis. For low EBV infection and normal vitamin D and estrogen levels we observe a healthy response, while for increased EBV or decreased vitamin D and estrogen, a disease outbreak is more likely. While the Level 2 model allows us to investigate the impact of potential risk factors on inflammation, it omits the impact of inflammation on demyelination and oligodendrocyte death.

**Figure 10:**
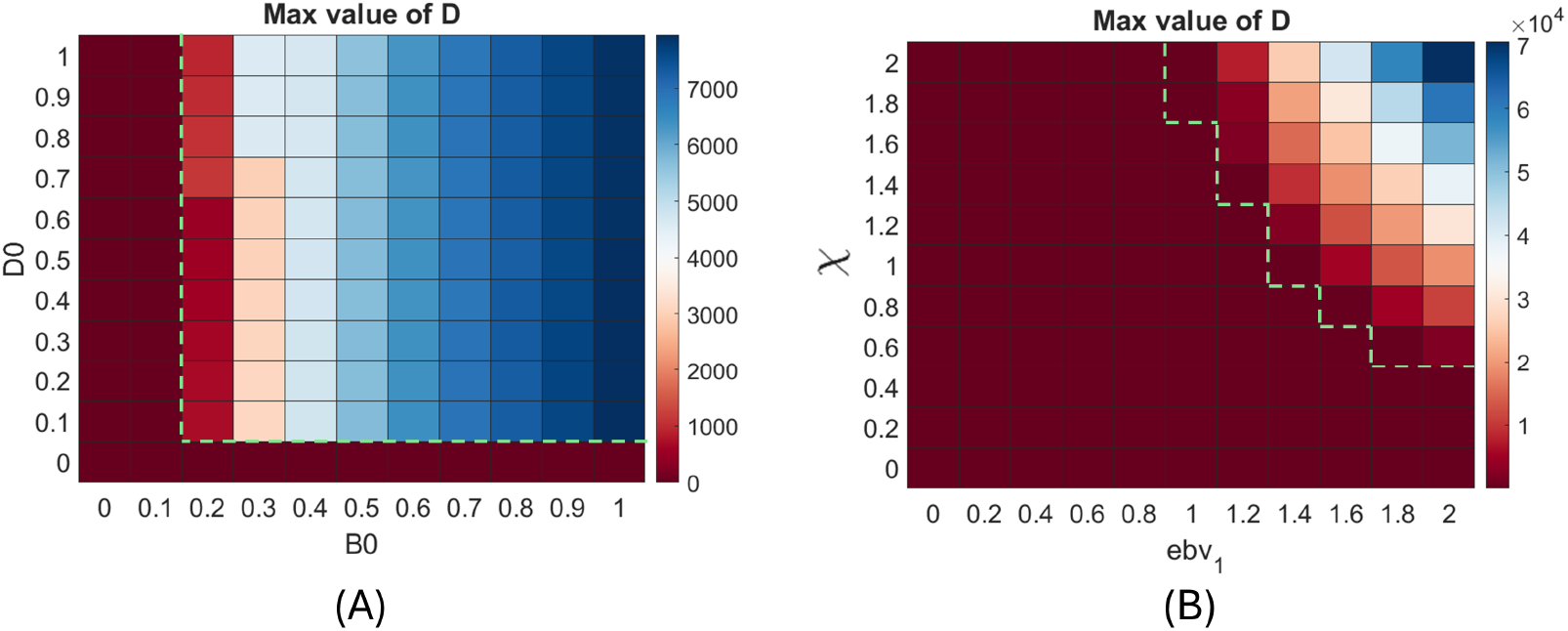
Heatmaps summarising the effects of changes to the initial concentration of (A) memory B cells *B*(0) and APCs *D*(0), and (B) estrogen *χ* and EBV *ebv*_1_, on the maximum value of APCs, i.e. *D*, from *t* = 0 to *t* = 200. The presence of oscillations is denoted by a green dashed line, where parameter values to the right of this line give rise to oscillations. The presence of oscillations was based on the criteria that three peaks were observed and the peaks were increasing in amplitude.

## 4. MS Model Level 3: Oligodendrocytes and myelin

A crucial aspect of MS disease is the demyelination of neurons, largely driven by effector immune cells, such as T cells, macrophages and B cells [8, 6], see Figure 1C. A natural extension of the Level 2 model is, therefore, to examine the impact of the current immune response on the oligodendrocytes and myelin in the CNS. To do this, we develop the Level 3 model, which introduces a population of effector immune cells whose response is linked to the immune cells in the Level 2 model, and populations of myelin and oligodendrocytes.

### 4.1. Level 3 model

To create the Level 3 model, we extended Equations (6)-(9) to include effector cells *E*(*t*), oligodendrocytes *O*(*t*), and myelin *M* (*t*):

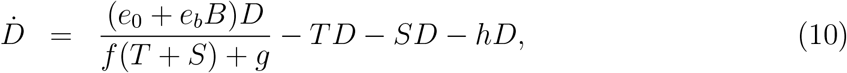

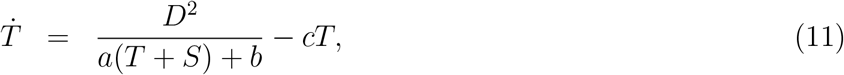

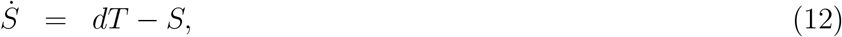

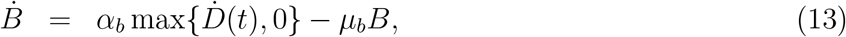

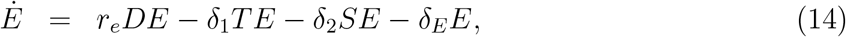

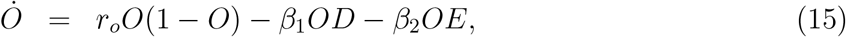

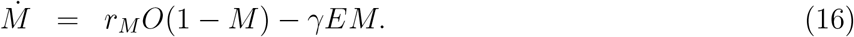

Here, effector cells are recruited by the presence of APCs at a rate *r*_*e*_ and regulated, i.e. inactivated, by regulatory cells at a rate *δ*_1_ and *δ*_2_. Oligodendrocytes divide logistically at a rate *r*_*O*_, and die from interacting with APCs or effector cells at a rate *β*_1_ and *β*_2_. Lastly, neurons are demyelinated at a rate *γ* by effector cells and regenerated by oligodendrocytes at a logistic rate *r*_*M*_ . The flow diagram for this model is given in Figure 11. We consider *O* and *M* to represent the oligodendrocytes and myelin in a small area of the brain as opposed to the whole brain.

**Figure 11:**
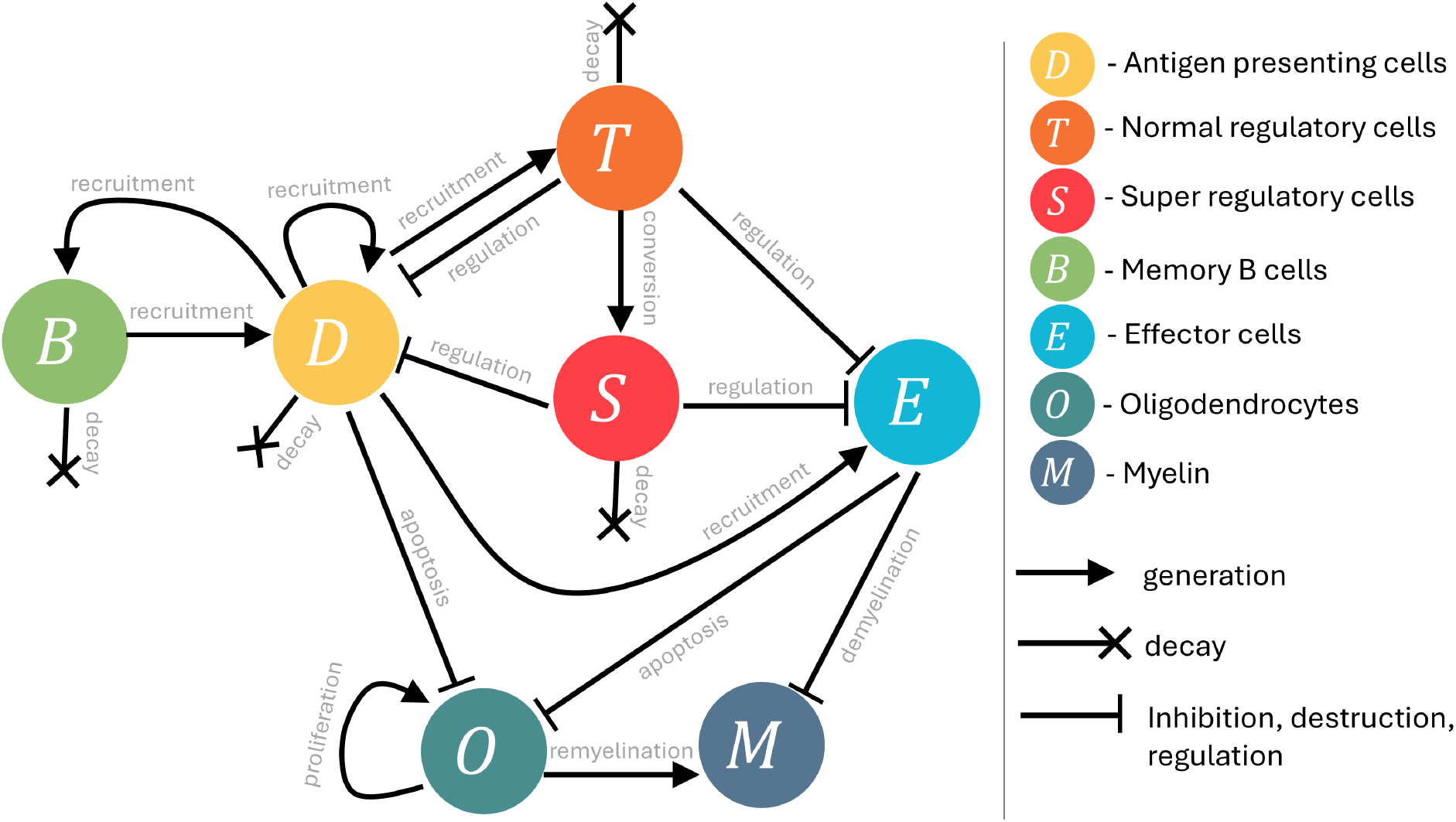
Level 3 model flow diagram corresponding to Equations (10)-(16). At this level, the focus is on incorporating effector immune cells and the effect of the immune response on myelin and oligodendrocytes. In this model, effector cells *E*, myelin *M*, and oligodendrocytes *O*, have been added to the previous Level 2 model. Effector cells are recruited by APCs and inhibited by regulatory cells. Effector cells and APCs caused damage to oligodendrocytes. Effector cells also cause damage to myelin. Oligodendrocytes regenerate and also regenerate myelin. Arrows represent promotion, flat bars represent inhibition, and crosses represent decay.

### 4.2. Model simulations

Given the detailed analysis on the possible link between inflammation and MS disease dynamics in the Level 1 and Level 2 models, we investigate whether these would translate to demyelination and oligodendrocyte damage in a full model of MS through numerical simulations. We simulate the model using the base parameterisation in Table 1. We chose *B*(0) = 0.01 as this was the initial condition considered for the Level 2 model that gave peripheral tolerance (see Figure 8). We discuss the parameter choices in Section 5. We make a number of observations:

#### Peripheral tolerance

We simulated the Level 3 model with the base parameters, see Figure 12A. Unsurprisingly, we see peripheral tolerance in the immune dynamics, i.e. *D* and *E*, leading to oligodendrocyte and myelin homeostasis, i.e. both populations, *O*(*t*) and *M* (*t*), returning to a value of 1. While Figure 12 shows the dynamics of *D*(*t*), *E*(*t*), *O*(*t*) and *M* (*t*), the other variables *T* (*t*), *S*(*t*) and *B*(*t*) are shown in the appendix in Figure A.13.

**Figure 12:**
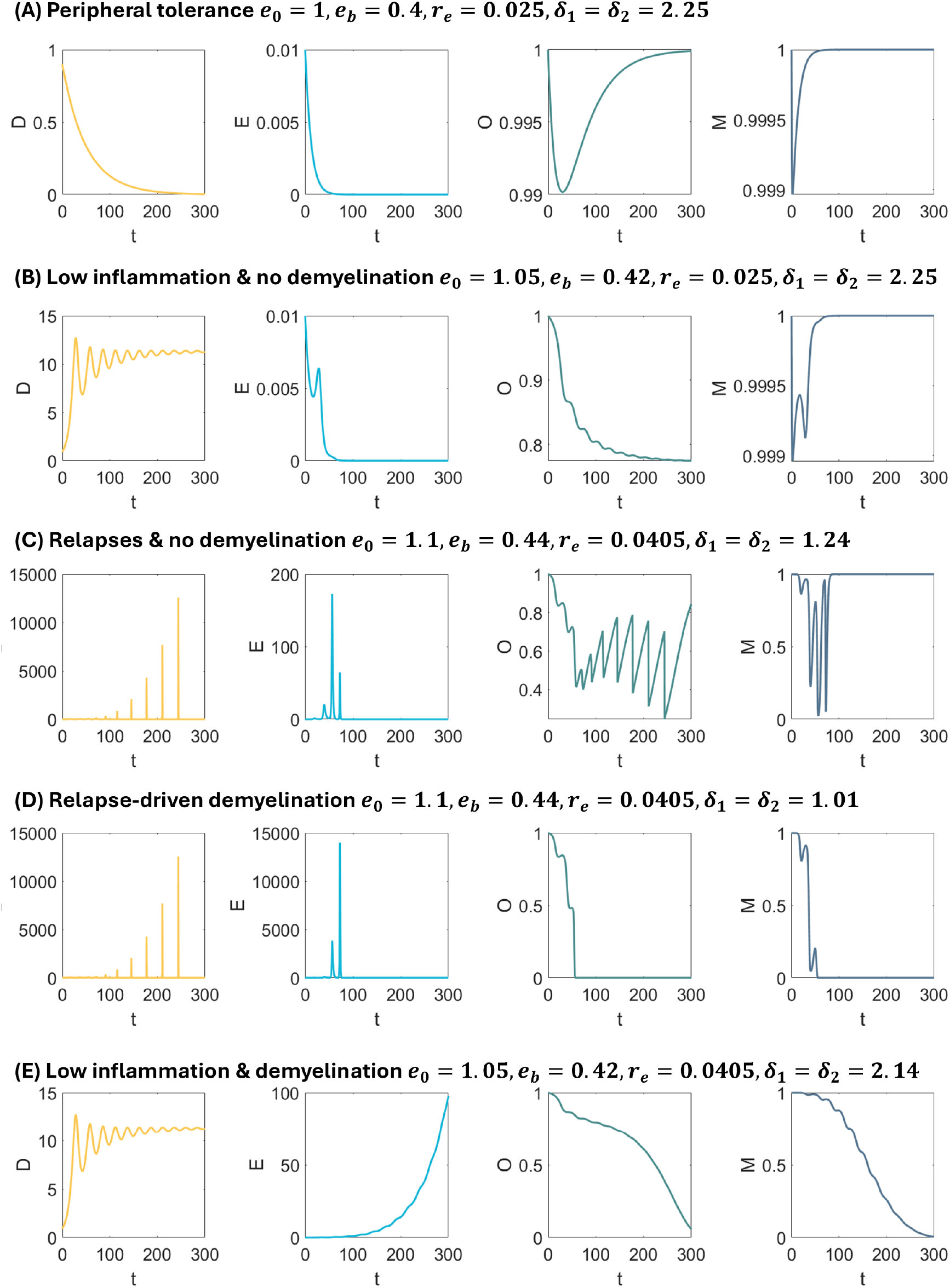
Level 3 model simulations for Equations (10)-(16). The model is simulated for the parameters in Table 1 and changes to any of these parameters are given in the figure titles. Specifically, parameters *e*_0_, *e*_*b*_, *r*_*e*_, *δ*_1_ and *δ*_2_ were changed across the subplots. The predicted dynamics for *D, E, O* and *M* are plotted with corresponding dynamics for *S, T* and *B* given in the Appendix.

#### Low inflammation & no demyelination

Following this, we focused our numerical simulations on examining the effect of small perturbations in parameters on the evolution of MS disease. We started by increasing the bifurcation parameter *e* that was analysed in the Level 1 model, see Figure 6. Increasing *e* corresponds to an increase in the APC generation. In the Level 2 model, *e* = *e*_0_ + *e*_*b*_*B*, so we increased *e*_0_ and *e*_*b*_ by 5% to examine the effect on the effector cells *E*, myelin *M* and oligodendrocytes *O* when the background immune response is at a non-zero, small steady state, see Figure 12B. We see that increasing *e*_0_ and *e*_*b*_ is enough to show a small decline in oligodendrocytes *O*, which stabilises as the APC population *D* stabilises. We also note that the effector population *E* declines back to zero and this allows the myelin *M* to fully recover. We denote this disease dynamic as ‘myelin resilience’, where we expect MS symptoms most likely would not be severe and the individual may not exhibit any MS disease markers.

#### Relapses & no demyelination

Continuing our investigation on the effect of increasing the APC generation through increasing *e*_0_ and *e*_*b*_, we increased their value from baseline by 10%. We also increased the arrival rate of effector cells *r*_*e*_ and decreased the clearance of effector cells, i.e. decreased *δ*_1_ and *δ*_2_. In doing this, we wanted to see the effect of a stronger effector response. We see in Figure 12C an even stronger immune response and increased relapses, however, this does not result in an increase in demyelination. At first this may be counterintuitive, as the hypothesis we were working with for the Level 1 and 2 models was that increased inflammation leads to increased demyelination and lesion generation. However, the model suggests that increased inflammation and relapses can lead to a down-regulation of the effector response as the regulatory response is so strong, allowing the myelin to recover. This suggests that the strength of relapses in a patient is not necessarily correlated to the amount of neurological damage. Future work could be done to investigate this more in-depth and to see how robust this is to perturbations.

#### Relapse-driven demyelination

Holding *e*_0_ and *e*_*b*_ at 10% of their baseline value, and decreasing the regulation of the effector cells, i.e. *δ*_1_ and *δ*_2_, results in a severe decline in the oligodendrocytes and myelin, Figure 12D. We see that if the effector cell population is more heavily dependent on the strength of the relapse, this can cause large numbers of effector cells that quickly deplete the myelin and oligodendrocytes. Interestingly, it is possible to obtain the same level of demyelination without strong relapses, but with changes to the effector cell population.

#### Low inflammation & demyelination

Taking the parameterisation of Figure 12B, if we also decrease slightly the strength of the regulatory response on the effector cells, i.e. *δ*_1_ and *δ*_2_, we see a very rapid demyelination, Figure 12E. This suggests that it is not necessarily the amount of effector cells that drives demyelination and oligodendrocyte death, but the ability of these cells to cause damage. Therapies that target reducing effector cell numbers may, therefore, not be effective for patients experiencing demyelination as in Figure 12E, as their effector cell numbers are already low. This could be an explanation for the varied efficacy of disease modifying therapies (DMTs), which look to decrease inflammation.

## 5. Parameter choices

The values of our base parameters are given in Table 1 and have been chosen as follows.

### Level 1

The parameter values for the Level 1 model are taken from [23], as listed at the top of page 2017 in [23]. The only difference is the choice of *d* = 4, which was *d* = 2 in [23]. We noticed that the length of the gap between signal bursts depends on *d*, and we increased *d* to get longer gaps. The parameter choice in [23] is based on a previous paper of the same group [53]. There it is assumed that the death rate of antigen presenting cells, effector T-cells and regulatory T-cells is approximately similar and given as 0.2 day^−1^. The death rate for the super regulatory cells is doubled, since they live less long. The decay rate of circulating antigen is assumed to be 5.6 day^−1^. The contact rate between T cells and APC is assumed to be 0.2 day^−1^ and about 1000 new T cells are generated from an antigen presentation event. Note that the parameters in Table 1 are based on a nondimensionalisation and a quasi-steady state assumption in [23], hence they are really combined parameters. Nevertheless, a time unit in our model corresponds to 5 days in reality.

### Level 2

Unfortunately, there is no data available to estimate parameters for models on Level 2 and Level 3; hence we chose parameters that we feel are biologically reasonable. The creation of memory *B*-cells is a slow process, much slower than the presentation of antigen. Hence here we assume the production of memory *B*-cells is about 1/100 of the production of antigen, i.e. *α*_*b*_ = 0.01. The death rate for memory *B* cells is chosen very small, *µ*_*b*_ = 0.0001, which corresponds to a half-life time of 94 years. The choice of *e*_0_ = 1 as base-APC generation corresponds to the healthy case, which is then increased by a 40% contribution of *B* cells, i.e. *e*_*b*_ = 0.4.

### Level 3

We set the reduction of effector cells by regulatory cells, *δ*_1_ and *δ*_2_, equal to the reduction in APCs, i.e. *δ*_1_ = *δ*_2_ = *h*. We set the generation of effector cells by APCs to be low: 1 APC generates an effector cell approximately every 40 days, i.e. *r*_*e*_ = 0.025. Oligodendrocytes proliferate slowly, especially when compared to myelin, i.e. *r*_*M*_ *>> r*_*O*_ and we expect remyelination to be occurring daily, whereas oligodendrocyte reparation would be closer to monthly, hence we chose *r*_*M*_ = 1.2 and *r*_*O*_ = 0.05. Oligodendrocytes are resilient and so we chose a small value for the loss of oligodendrocyte due to the immune attack, i.e. *β*_1_ = *β*_2_ = 0.001. In comparison, we expect the demyelination of neurons by effector cells to be faster than the oligodendrocyte loss, i.e. *γ > β*_1_ = *β*_2_, and so we chose *γ* = 0.15. There are other biologically reasonable values for these parameters, and it is possible that *δ*_1_ ≠ *δ*_2_ and *β*_1_ ≠ *β*_2_; however, we felt a full exploration of the parameters is beyond the scope of this work.

## 6. Discussion and Conclusion

Multiple sclerosis (MS) is an inflammatory disease of the brain and spine, where the immune system damages protective myelin coating of neurons and causes significant physical and cognitive decline. The incidence of MS is increasing, particularly in women, demonstrating a paramount need for reliable mathematical modelling that can be used to interrogate the many unknowns in the disease. In this work, we propose a new paradigm for mathematical modelling of MS focusing on three levels capturing self-antigen presentation and relapsing-remitting MS (Level 1), peripheral tolerance and immune memory (Level 2) and damage to myelin and oligodendrocytes (Level 3).

In this work, we were motivated by a list of key questions underpinning MS disease dynamics that we felt a modelling paradigm should be able to answer (see Figure 3). Considering only the interaction between APCs and regulatory T cells in the Level 1 model, we were able to find parameter conditions under which we could describe short outbreaks in inflammation followed by long transient phases, such as those seen in RRMS (Q3). We found that increasing the recruitment rate of APCs, *e*, would lead to oscillations of heightening amplitude, reminiscent of RRMS. We felt this was reminiscent of the effects of estrogen on the immune system, as estrogen is known to sensitize the immune system to self-antigen presentations (Q7).

Introducing memory B cells into the Level 1 model, we further investigated the impact of estrogen on disease dynamics (Q7) with the Level 2 model. We found that both low estrogen and low vitamin D had similar effects and led to long durations of intermediate oscillations. Most importantly, the addition of memory B cells introduced stable peripheral tolerance into the model dependent on the initial number of memory B cells. This suggests that the generation of a large memory B cell pool may be a driving factor in disease onset.

Finally, we extended the Level 2 model to include effector immune cells and their effect on myelin and oligodendrocytes. Interestingly, we see more complicated dynamics arising from the introduction of the new populations, with both relapses and effector cells being possible sources of demyelination in the CNS. We leave it for future work to conduct a complete analysis of the Level 3 model, and present it as a possible simple way to incorporate effector cells and the oligodendrocyte/myelin population into a model of MS disease.

We believe our modelling can contribute to the discussion on the MS prodrome [15]. The stable coexistence steady state that arose in our analysis carries all the features of a healthy-like state, which is not clinically detected, but indicates already a first sign of trouble. This steady state should lead to increased neurofilament light chain (NfL) concentrations in plasma, which could possibly be identified if repeated NfL measurements are raised and constant over time. This could be an interesting thought for further exploration.

One major limitation of our framework is the lack of calibration to data. Unfortunately, it is challenging, and in some instances impossible, to calibrate the parameters in the model to data. Real human data at the cellular level is not possible to obtain and current imaging techniques are not refined enough to capture the microscale dynamics in the brain. There is data from mouse models called the Experimental Autoimmune Encephalomyelitis (EAE) model; however, this model is thought to differ from the human MS disease. For now, we leave these parameters as undetermined but reasonable values and conduct a more theoretical exploration.

We believe that the hesitancy to model MS with a mathematical approach stems from the high level of complexity of the disease. Not only are many different types of immune cells involved, the disease also progresses through multiple phases with very different characteristics. In addition, MS is expressed quite differently from patient to patient. We think the systematic classification of the underlying mechanisms into Level 1-6 can form a basis for discussions. Models can be developed on each level, and possibly experiments on each level can be devised to help estimate model parameters.

In summary, the mathematical models presented here are able to capture a range of MS disease dynamics from peripheral tolerance, to relapses and progressive disease. A natural future extension of this work would be to consider the effects of treatment. There are nine classes of disease-modifying therapies (DMTs) approved for treatment of RRMS or SPMS [54]. The efficacy of current DMTs varies significantly from 29% to 68%, highlighting a crucial need to understand their role in the immunopathology of MS using a mathematical model [54]. With small modifications to the model capturing treatment pharmacokinetics and pharmacodynamics, it would be possible to investigate the efficacy of DMTs with our Level 2 and 3 models. We believe that future work could build off our framework and develop useful tools for the diagnosis and treatment of MS disease.

## Appendix A. Level 3 model additional simulations

We first note that the Level 3 model contains a steady state solution of peripheral tolerance, i.e.

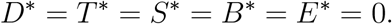

and

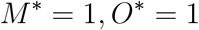

We also note, that if we assume *E* and *D* are at steady state, i.e. *E*^∗^ and *D*^∗^, then the steady state values for *O*^∗^ and *M* ^∗^ are written as

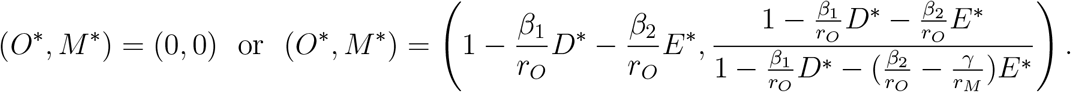

This suggests that if biologically reasonable parameter values exist for which either of these steady states is stable and non-zero, we can achieve peripheral tolerance and chronic inflammation with the Level 3 model. This is evidenced by the simulations obtained in Figure 12A and B.

The simulations for *S, T* and *B* corresponding to the simulations in Figure 12 are given in Figure A.13.

**Figure A.13:**
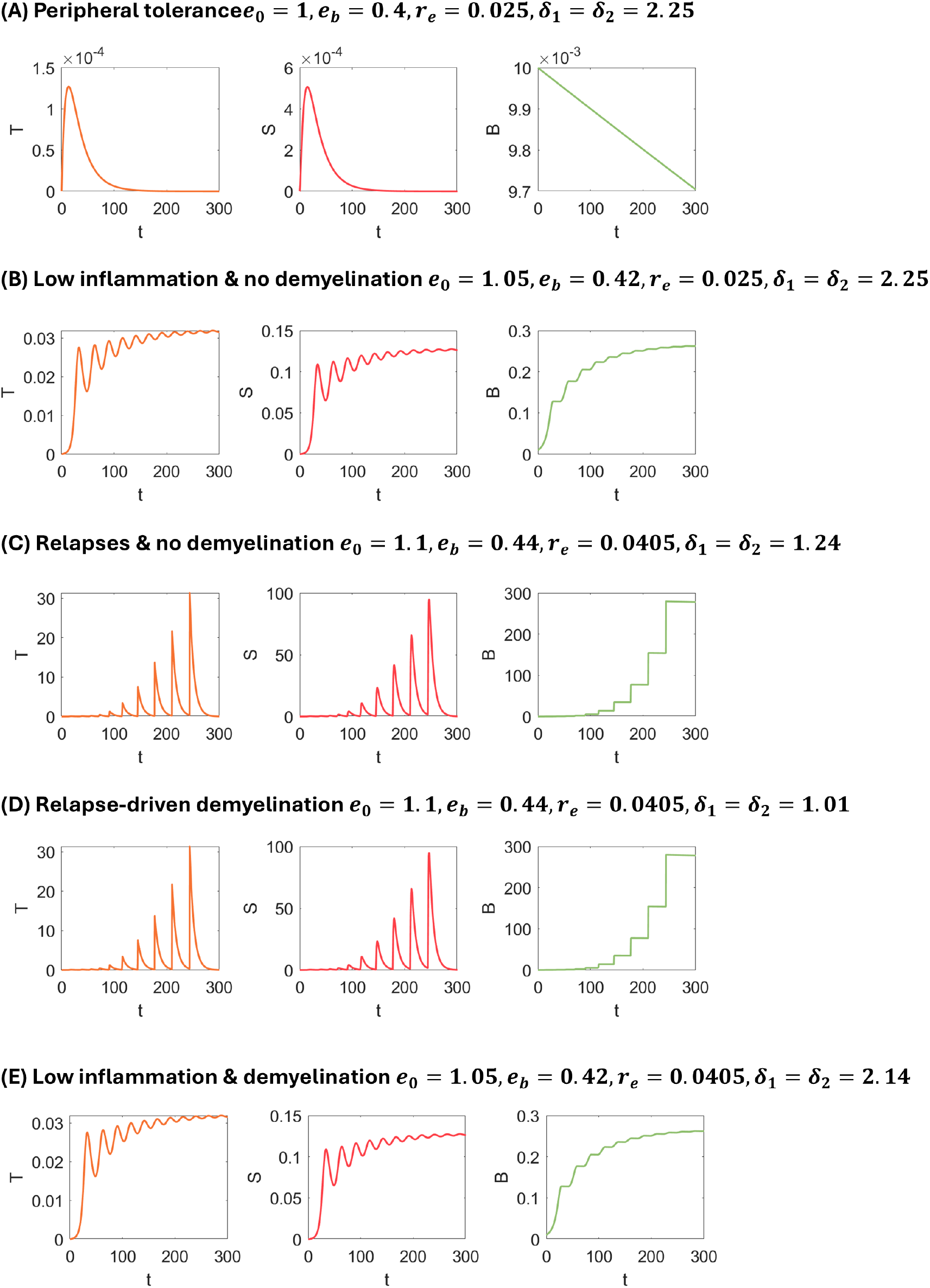
Level 3 model simulations for Equations (10)-(16) corresponding to Figure 12. The model is simulated for the parameters in Table 1 and changes to any of these parameters are given in the figure titles.

